# High sugar diets can increase susceptibility to bacterial infection in *Drosophila melanogaster*

**DOI:** 10.1101/2023.12.07.570705

**Authors:** Andrea M. Darby, Destiny O. Okoro, Sophia Aredas, Ashley M. Frank, William H. Pearson, Marc S. Dionne, Brian P. Lazzaro

**Affiliations:** Department of Entomology, Cornell University, Ithaca, NY, United States of America; Cornell Institute of Host-Microbe Interactions and Disease, Cornell University, Ithaca, NY, United States of America; University of California, Irvine, Irvine, CA, United States of America; Department of Microbiology, Cornell University, Ithaca, NY, United States of America; Battelle, Columbus, Ohio, United States of America; Department of Life Sciences, Imperial College London, London SW7 2AZ, United Kingdom; Centre for Bacterial Resistance Biology, Imperial College London, London SW7 2AZ, United Kingdom

**Keywords:** *Drosophila*, high-sugar, innate immunity, metabolism, bacterial infection

## Abstract

Overnutrition with dietary sugar can worsen infection outcomes in diverse organisms including insects and humans, generally through unknown mechanisms. In the present study, we show that adult *Drosophila melanogaster* fed high-sugar diets became more susceptible to infection by the Gram-negative bacteria *Providencia rettgeri* and *Serratia marcescens,* although diet had no significant effect on infection by Gram-positive bacteria *Enterococcus faecalis* or *Lactococcus lactis.* We found that *P. rettgeri* and *S. marcescens* proliferate more rapidly in *D. melanogaster* fed a high-sugar diet, resulting in increased probability of host death. *D. melanogaster* become hyperglycemic on the high-sugar diet, and we find evidence that the extra carbon availability may promote *S. marcescens* growth within the host. However, we found no evidence that increased carbon availability directly supports greater *P. rettgeri* growth. *D. melanogaster* on both diets fully induce transcription of antimicrobial peptide (AMP) genes in response to infection, but *D. melanogaster* provided with high-sugar diets show reduced production of AMP protein. Thus, overnutrition with dietary sugar may impair host immunity at the level of AMP translation. Our results demonstrate that dietary sugar can shape infection dynamics by impacting both host and pathogen, depending on the nutritional requirements of the pathogen and by altering the physiological capacity of the host to sustain an immune response.

**Author Summary:** Diet has critical impact on the quality of immune defense, and high-sugar diets increase susceptibility to bacterial infection in many animals. Yet it is unknown which aspects of host and pathogen physiology are impacted by diet to influence infection dynamics. Here we show that high-sugar diets increase susceptibility to some, but not all, bacterial infections in *Drosophila*. We find that feeding on high sugar diet impairs the host immune response by reducing the level of antimicrobial peptides produced. The expression of genes encoding these peptides is not affected, so we infer that protein translation is impaired. We further show that flies on high-sugar diets are hyperglycemic, and that some pathogens may use the excess sugar in the host to promote growth during the infection. Thus, our study demonstrates that dietary impacts on infection outcome arise through physiological effects on both the host and pathogen.

## Introduction

Nutritional status and diet are important factors for immune defense in both mammals and insects (1–5). Hyperglycemia and diets excessively high in sugar have adverse effects on metabolic homeostasis and infection outcome (6–10). For example, human patients admitted to the intensive care unit with sepsis have a higher probability of death if they have elevated blood sugar (11) and non-diabetic hyperglycemia increases the severity of *Mycobacterium tuberculosis* infections in guinea pigs (1). A similar phenomenon is observed in the fruit fly *Drosophila melanogaster*, where adults reared on high sugar diets experience higher pathogen burden (3) and increased mortality from systemic bacterial infection (2,12). Despite these clear effects of diet and nutritional state on infection outcome, the molecular and physiological mechanisms by which high sugar impacts infections is often unclear. By uncovering these mechanisms, we can clarify the role that diet has on shaping organismal physiology during an active infection and identify strategies to alleviate consequences of infection that are exacerbated by obesogenic diets.

*Drosophila* is an excellent model to understand the mechanisms by which high-sugar diets affect infection dynamics, with extensive prior study on infection, diet, and metabolism (e.g., 13– 16). Lacking adaptive immune systems, *Drosophila* rely on innate immune defenses that include humoral production and secretion into circulation of antimicrobial peptides (AMP) and cytokines secreted that share homologous function with innate immune defenses in mammals and other invertebrate species (17,18). The Toll and the immune deficiency (IMD) pathways are the two major regulators of the *Drosophila* humoral immune response to infection (14). Gram-positive bacterial and fungal infections predominantly stimulate Toll activity (19,20) while Gram-negative bacterial infections activate the IMD pathway (19,21). Once activated, both pathways lead to the nuclear translocation of nuclear factor-κB (NF-κB) family transcription factors to drive expression of hundreds of infection-responsive genes, including those encoding antimicrobial peptides (22,23). Infection and/or constitutive activation of these pathways stimulates major shifts in metabolic processes to support the immune response, including suppressed insulin signaling, reduction in energetic stores like glycogen and triglycerides, and alterations in carbohydrate and lipid metabolism (24–28).

Both high-sugar diets and infection alter metabolism and energetic usage in *Drosophila* (29). In the absence of infection, high-sugar diets can also upregulate genes involved in the immune response, including antimicrobial peptides (12,30), which is similar to high levels of dietary sugar stimulating low-grade inflammation responses in humans (31). Additionally, uninfected *Drosophila* larvae reared on high-sugar diets exhibit impaired melanization and reduced phagocytic capacity of fungal spores (12,32). While there is evidence that high-sugar diets affect the *Drosophila* immune response in uninfected states, it is less clear whether high-sugar diets impact immune activity during a live infection. Resistance to infection can be measured as immune system activity or control of pathogen burden (33). Feeding *Drosophila* high-glucose diets results in higher pathogen burden after systemic infection with the Gram-negative bacteria *Providencia rettgeri* (2,3) and *Pseudomonas aeruginosa* (6). This could suggest that high-sugar diets impair host immune system activity, but *Pseudomonas aeruginosa* is additionally able to use glucose in hyperglycemic hosts to establish higher pathogen burdens (6). Thus, it has been difficult to discern whether elevated pathogen burden in *Drosophila* provided with high-sugar diets is due to impaired host immune responses or to bolstered pathogen growth capacity. These are not mutually exclusive alternatives. Furthermore, prior *D. melanogaster* studies that investigated the effects of high dietary sugar on systemic infection outcome were performed using flies that were reared throughout their entire development on high-sugar diets (2,3,12,32). Rearing on elevated dietary sugar diets causes hyperglycemia, lipidemia, and reduced body weight in both larvae and as adults, and prolongs larval developmental time (12,34). Thus, it is difficult to determine in those experiments whether effects of diet on adult infection outcome result from altered metabolism and immunity or from indirect consequences of altered development.

In the present study, we specifically focus on the immediate effects of high dietary sugar on infection dynamics in adult *D. melanogaster*. We do this by rearing *D. melanogaster* larvae on a common base diet, then switching adults to diets that vary in sugar content prior to delivering systemic bacterial infection. To determine whether adverse effects from high-sugar diets are universal across pathogenic infections, we test infection with four different bacterial pathogens representing Gram-positive and Gram-negative bacteria. We find that flies provided with high-sugar diet as adults are particularly susceptible to Gram-negative bacterial infection, suffering increased mortality and more rapid pathogen proliferation during the early hours of infection. We find evidence that increased carbon availability in high-sugar hosts can accelerate the growth of certain pathogens, and that high-sugar diets impaired immune system function at the level of antimicrobial peptide translation without affecting transcription levels.

## Results

### High-sucrose diets increase mortality from infection with some bacterial pathogens

We first wanted to test whether high-sugar diets reduce immune defense in adult *D. melanogaster* independent of any developmental consequences. We therefore reared larvae on a yeast-cornmeal diet containing 4% sucrose, then split the population across the experimental diets during the non-feeding pupal stage such that eclosing adults emerged onto experimental diets that were 0%, 2%, 4%, 8%, 16%, or 24% (w/v) sucrose (Fig. 1A), which covers the range used in prior studies (3,12,32). After 3-5 days on the experimental diets, adult female flies were given a systemic infection with one of four bacteria: *Providencia rettgeri*, *Serratia marcescens, Enterococcus faecalis, Lactococcus lactis*. These bacteria were chosen as natural pathogens of *D. melanogaster* (35,36) and include both Gram-positive and Gram-negative bacteria, which are respectively expected to predominantly activate the Toll and IMD immune response pathways (19).

**Fig 1.**
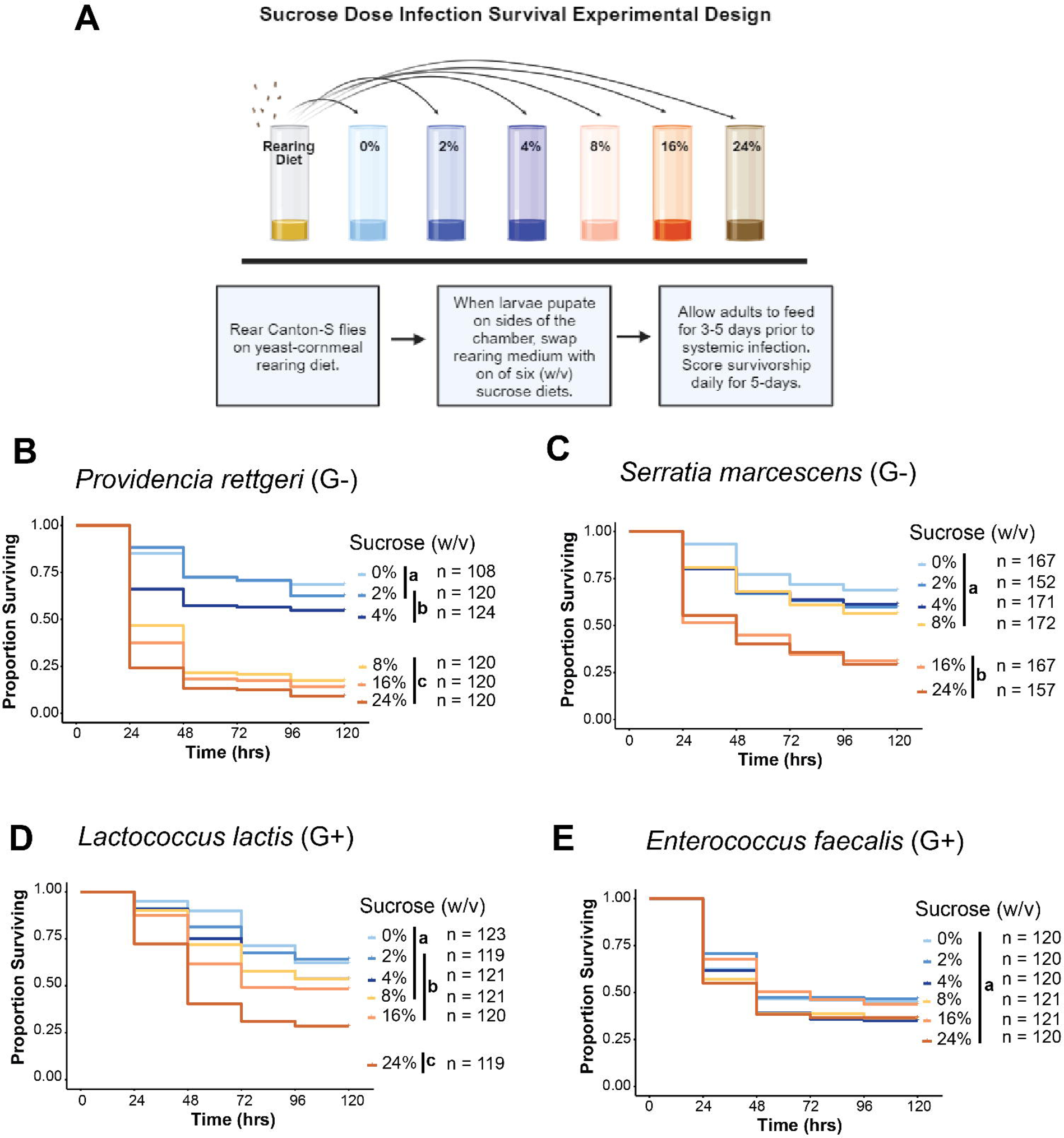
High-sugar diet increases mortality after infection with some pathogens. (A) Experimental setup for assaying effects of high-sugar diets on adult survival of infection. The six experimental diets used have the same yeast and cornmeal content, but they varied in sucrose (w/v). Diets with 0%, 2%, and 4% are represented in shades of blue while diets with 8%, 16%, 24% sucrose are represented in shades of orange. The rearing diet contains 4% sucrose. (B) Flies fed 8%, 16%, and 24% sucrose diets exhibit significantly higher mortality after *Providencia rettgeri* infection than flies given 0%, 2%, and 4% sucrose diets (p<0.0001; Cox Mixed Effects Model). There was no difference in survival among flies provided with the three high-sugar diets, while flies provided with the 4% sucrose diet exhibited significantly lower survivorship than flies provided with 0% added sucrose (p = 0.035; Cox Mixed Effects Model). (C) Flies fed 16% and 24% sucrose diets exhibit significantly higher mortality after *Serratia marcescens* than flies given diets with 8% sucrose or lower (p<0.0001; Cox Mixed Effects Model). (D) Flies fed the 24% sucrose diet have the highest mortality after *Lactococcus lactis* infection compared to all other diets (p<0.001, Cox Mixed Effects Model). Flies fed 16% sucrose died significantly faster than flies fed 0% sucrose diets (p = 0.047; Cox Mixed Effects Model) but there were no other significant pairwise differences in survivorship among flies provided the 16% sucrose diet or lower. (E) There was no effect of diet on survivorship after infection with *Enterococcus faecalis*. Letters denote significant pairwise differences between diets (p<0.05).

**Fig. S1.**
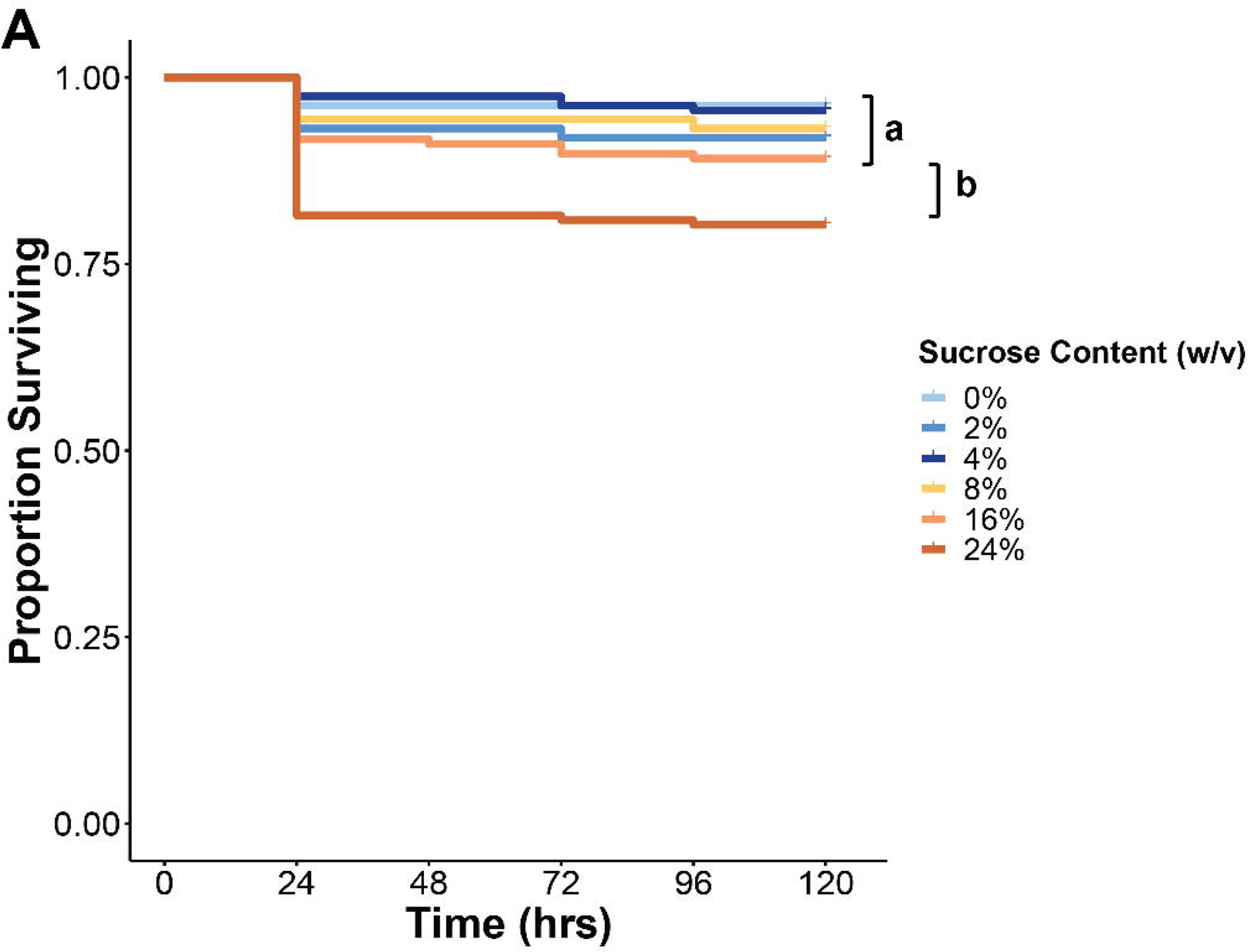
A) Survivorship of PBS injury controls across all infection experiments pooled together. Flies fed 24% (w/v) sucrose diet have higher proportion of death after sham-infection with sterile PBS compared to all other diets (p<0.05; Cox Proportional Hazards Mixed effects model).

**Fig. S2.**
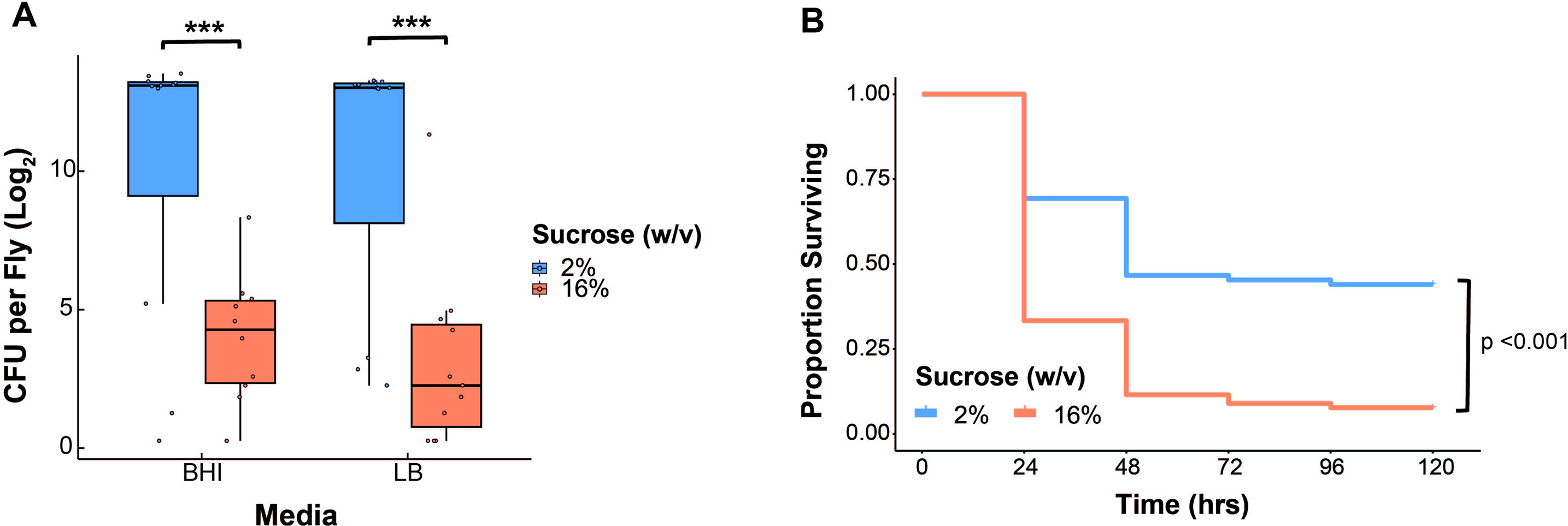
A) To measure the abundance of the endogenous gut microbiota in uninfected flies fed on the 2% (w/v) sucrose or 16% sucrose diets, 10 whole flies were pooled and homogenized in 250 μl of PBS, and 50 μl of the homogenate was plated on lysogeny broth (LB) or brain heart infusion (BHI) agar. Flies fed on the 2% diets exhibit higher microbiota loads on LB (p<0.001, n = 15 pools of 10 flies per diet) and BHI (p<0.001, n = 15 pools of 10 flies per diet) plates than flies fed 16% sucrose. C) Axenic flies were infected with *P. rettgeri*, and flies fed 16% sucrose diets exhibited higher mortality than flies given 2% sucrose (p<0.001; Cox Proportional Hazards model).

We found that flies provided with high-sucrose diets experienced higher mortality after infection with the Gram-negative bacteria *S. marcescens* and *P. rettgeri* than flies fed low sugar (Figs. 1B and 1C). This increased mortality occurred in a concentration-dependent manner depending on the pathogen. Flies provided with 0%, 2%, or 4% sucrose experienced low mortality and flies provided with 16% or 24% sucrose experienced high mortality after infection with both pathogens. However, on the 8% sucrose diet, flies infected with *S*. *marcescens* exhibited high survival (Fig. 1B), but flies infected with *P. rettgeri* exhibited low survival (Fig. 1C). Overall, there was no effect of diet on survivorship of infection with the Gram-positive bacteria *L. lactis* and *E. faecalis* (Figs. 1D and 1E). Although flies provided with the 24% sucrose diet experienced the highest mortality after *L. lactis* infection (Fig. 1D), we attribute this to the diet being generally poor as sham-infected control flies fed on the 24% sucrose diet also experienced elevated mortality (Fig. S1A). Varying the diet of the host can alter the composition of the gut microbiota (37–39), which can yield effects on the host’s metabolic status (40–42). Flies provided with 16% sucrose in our experiments showed a reduced abundance of gut flora compared to flies provided with 2% sucrose diet (Fig S2A). However, we found that high-sugar diets increased the probability of death from systemic *P. rettgeri* infection even in axenic flies with no microbiota (Fig S2B), suggesting that our observed effect of diet on immunity is not mediated by effects on gut flora.

### Flies fed on high-sucrose diets become hyperglycemic despite eating less

*D. melanogaster* will modulate feeding behavior depending on the content of their diet (43–45), and the nutritional composition of the diet influences metabolic state (3,34,46). We therefore first assessed whether varying sucrose levels in the diet impacts feeding rate. We performed an excreta quantification assay (ExQ) (47) on uninfected females to measure feeding across the six experimental diets shown in Figure 1A. Flies were housed in ExQ chambers over a 2-day period with experimental diets containing the dye erioglaucine (1% w/v) and their excreta was collected every 24-hours. We found that flies ate significantly more of the three diets with the lowest sucrose (0%, 2%, and 4%) than the three diets with the highest sucrose (8%, 16%, 24%; p < 0.0001, Fig. 2A). There was no significant difference in feeding rate among the three lowest-sucrose diets or among the three highest-sucrose diets. In subsequent experiments, we focus on the 2% and 16% sucrose diets as representative the low-and high-sugar diets.

**Fig. 2.**
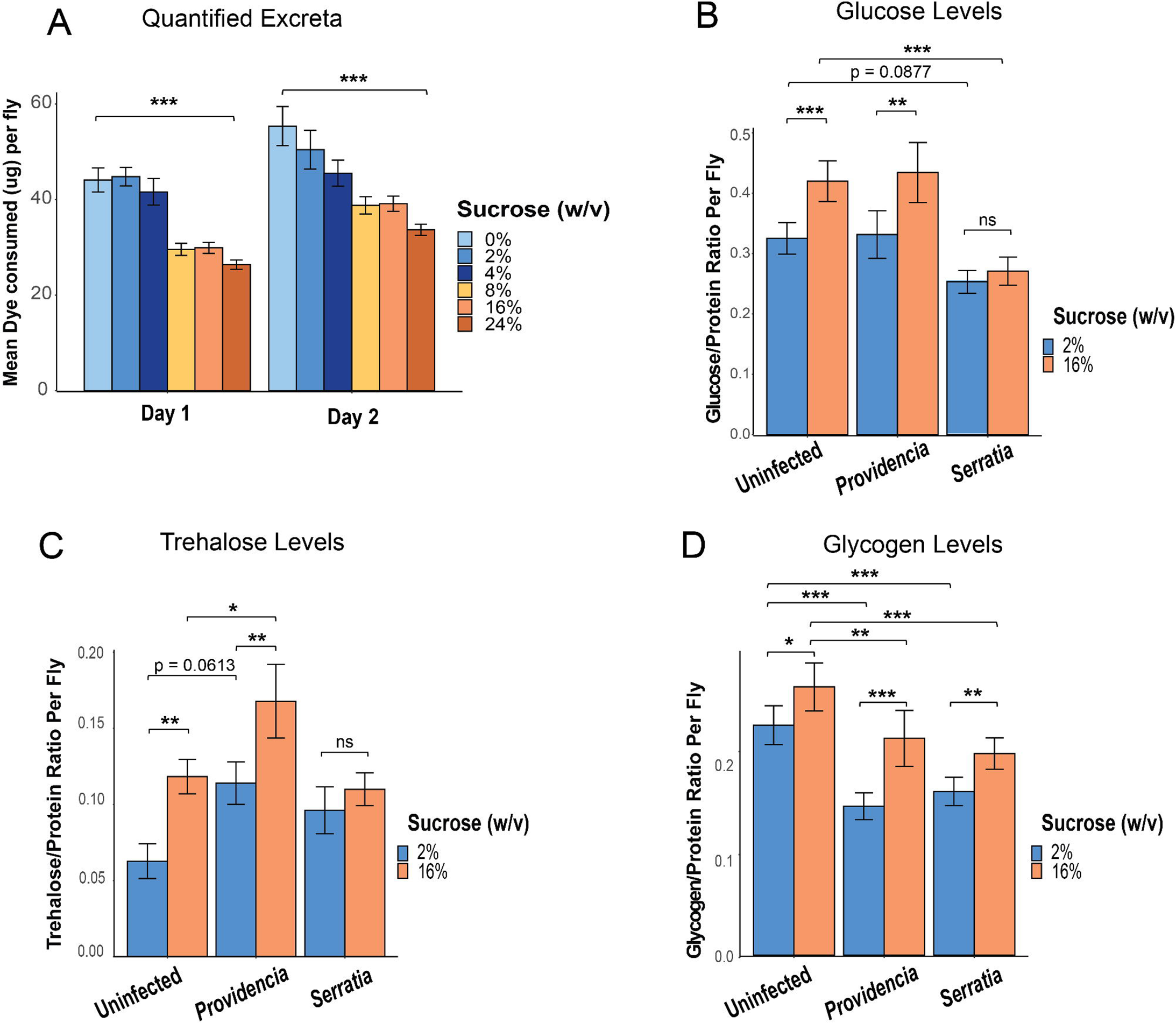
Flies fed high sugar sustain higher carbohydrate stores despite reduced feeding. (A) Excreta quantification of uninfected female flies fed on diets ranging from 0%-24% sucrose was measured over two days. Flies feeding on the 8%, 16%, and 24% diets consumed significantly lower dye than those fed on the 0%, 2%, 4% sucrose diets (p < 0.001). (B) Uninfected flies fed on the 16% diet had higher glucose levels compared to flies fed on 2% sucrose (p < 0.001; ANOVA with Tukey HSD post-hoc). After 24 hours of *P. rettgeri* infection, flies fed on high sugar continued to exhibit higher glucose levels compared to flies fed 2% sucrose (p < 0.01; ANOVA with Tukey HSD post-hoc). However, there was no difference in glucose levels between flies provided with 2% and 16% diets 24-hours after *S. marcescens* infection (p=0.658). Flies fed 16% sucrose exhibit significantly reduced glucose levels after *S. marcescens* infection (p<0.001), and there was a nearly significant decrease in glucose levels of flies fed 2% sucrose after *S. marcescens* infection (p = 0.0877) (C) Trehalose levels were significantly higher in flies fed 16% sucrose than in flies fed 2% sucrose when the flies were uninfected (p<0.001) or when they were infected with *P. rettgeri* (p<0.01). However, *S. marcescens* infection had no effect on trehalose levels between flies given the 2% and 16% diets (p = 0.38). *P. rettgeri* infection led to higher trehalose levels in flies on both 2% (p = 0.0554) and 16% (p = 0.0446) sucrose diets compared to uninfected flies, while *S. marcescens* had no effect on trehalose levels on either diet. (D) Glycogen levels were significantly higher in flies on 16% versus 2% diets when flies were uninfected (p = 0.0479), infected with *P. rettgeri* (p = 0.019), or infected with *S. marcescens* (p = 0.035). On both diets, infection with both pathogens led to reduced glycogen levels compared to uninfected (2% *Providencia*, p<0.01; 16% *Providencia,* p = 0.036; 2% *Serratia*, p<0.01; 16% *Serratia*, p < 0.01). Legend for panel figure: * = p<0.05, ** = p<0.01, *** = p <0.001.

To assess the effects of diet on the metabolic status of adults, we measured glucose, trehalose, glycogen, and soluble protein in the whole body of uninfected flies provided with either 2% or 16% sucrose diets. We found that uninfected flies fed the high-sucrose diet exhibit higher levels of glucose (p < 0.001, Fig. 2B), trehalose (p = 0.001, Fig. 2C), and glycogen (Fig. 2D, p= 0.0479) than flies fed the low-sugar diet (Fig. 2B-D). Given that infection can stimulate shifts in carbohydrate levels (26,48), we also tested whether flies fed high-sugar continue to exhibit elevated carbohydrate levels after infection. We indeed found that flies fed the high-sucrose diet exhibited higher levels of glucose (p<0.01, Fig. 2B), trehalose (p < 0.01, Fig. 2C), and glycogen (p =0.019, Fig. 2C) than flies fed the low-sucrose diet at 24 hours after *P. rettgeri* infection. Interestingly, there was no difference in glucose content (Fig. 2B) or trehalose content (Fig. 2C) between flies fed low-sucrose or high-sucrose at 24 hours after infection with *S. marcescens*. However, flies given a high-sucrose diet exhibited higher glycogen stores after *S. marcescens* infection than flies given low sucrose (p = 0.035 Fig. 2D). Despite reduced feeding on high sugar, flies on the high-sucrose diet still sustain higher levels of carbohydrate stores in both uninfected and infected states.

### Feeding on high sucrose increases pathogen load in the early stages of infection

High pathogen load is associated with higher mortality from infection (49,50). Since we observed that high-sugar diets lead to higher mortality during infection with the Gram-negative bacteria *S. marcescens* and *P. rettgeri*, we wanted to establish whether this high mortality arises from higher bacterial load. To test whether flies fed on high sugar have higher pathogen loads, we performed an *in vivo* bacterial growth assay where we sampled infected flies at two-hour intervals from 0-16 hours post-infection and at 24, 36, and 48 hours post-infection. At each time of sampling, individual flies were homogenized and plated on agar plates to estimate the number of live bacterial cells in the fly.

Overall, the bacteria proliferated more quickly in flies fed on the high-sugar diet after both *P. rettgeri* (Fig. 3A) and *S. marcescens* (Fig. 3B) infection. However, the divergence in load between the two diets occurs at different times for the two pathogens. *S. marcescens* infections start to exhibit a higher pathogen burden as soon as 6 hours into the infection, while *P. rettgeri* burden becomes higher by 12 hours into the infection. Flies that died during the sampling period were not included in this assay, which may mean that pathogen burdens are underestimated if the surviving flies at any given timepoint tend to be the ones with lower pathogen burdens. At 10 hours and later post-infection with *S. marcescens*, we observed a lower proportion of surviving flies on the high-sugar diet than on the low-sugar diet (Fig. 3B), which may explain the apparent absence of difference in pathogen burden 10-16 hours into the infection. We similarly observe a lower proportion of flies provided with high-sugar diet surviving *P. rettgeri* infection 10 hours and later, although the difference in pathogen burden between flies on the two diets remains significant (Fig. 3A).

**Fig 3.**
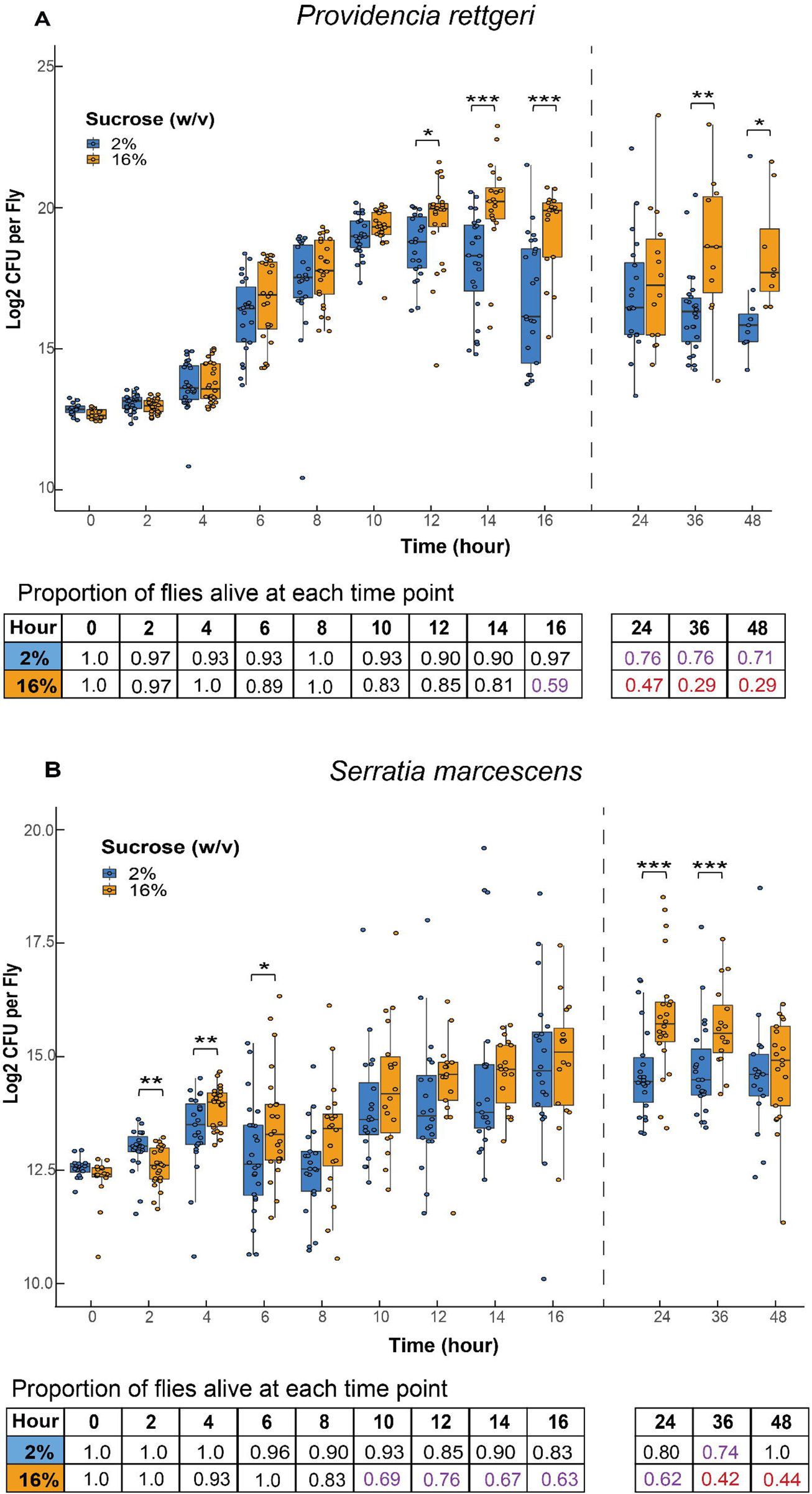
Pathogens proliferate faster in flies fed high sugar. (A) Pathogen load was assayed at 2-hour intervals from 0-16 hours post-infection and at 24, 36, and 48 hr post-infection in flies fed 2% and 16% sucrose diets. After *P*. *rettgeri* infection, flies provided with high-sugar diets have higher pathogen loads at 12, 14, 16, 36, and 48 hr than flies given low-sugar diet (12 hr, p =0.031; 14 hr, p <0.001, 16 hr, p< 0.001; 36 hr, p < 0.01; 48 hr, p = 0.022; Generalized Least Square Means (GLS) with Tukey HSD post-hoc for pairwise comparisons). Flies given high-sugar diets have consistently lower survival to time of sampling at hour 10 and onward. (B) After *S. marcescens* infection, flies given high-sugar diets have higher pathogen loads than flies given low-sugar diet at 4 hr, 6 hr, 24 hr, and 36 hr post-infection (4 hr, p = 0.011; 6 hr, p = 0.044, 24 hr, p< 0.001; 36 hr, p < 0.01; Generalized Least Square Means (GLS) with Tukey HSD post-hoc for pairwise comparisons). Flies given high-sugar diets have consistently lower survival to time of sampling at hour 8 and beyond. At 2 hours post-infection, flies given low-sugar diets have higher pathogen load than flies given high-sugar diets (p < 0.01). Each dot represents the pathogen burden of an individual fly. Legend for panel figure: * = p<0.05, ** = p<0.01, *** = p <0.001.

### Nutrient availability promotes *S. marcescens* proliferation within hosts on high-sugar diets

Having determined that *D. melanogaster* provided with high-sucrose diets are both more susceptible to infection and sustain higher internal carbohydrate stores, we next sought to understand whether the increased susceptibility to infection on the high-sugar diet is due to impaired immune system activity or to an altered nutritional environment experienced by the infecting pathogen. To test this, we measured pathogen proliferation during *in vivo* infection of immune compromised *D. melanogaster* on the low-and high-sugar diets. We predicted that the effect of diet on sensitivity to infection would be eliminated in immunocompromised flies if the dietary effect is due to impaired immunity, but it would remain if the dietary effect were due to altered nutritional environment for the pathogen. We used the *D. melanogaster* strain ΔAMP10 (51), which has intact Toll and IMD signaling but is missing 10 genes encoding major infection-inducible antimicrobial peptides. Antimicrobial peptides are small, cationic peptides that are secreted from the insect fat body in response to a systemic infection, and they are critical for killing pathogens by disrupting function of specific microbial processes (52–55). The ΔAMP10 flies have mutations that eliminate *Diptericin A*, which is critical for controlling *P. rettgeri* infection (51,56), and the four *Attacin* genes, three of which are critical for the control of *S. marcescens* infection (P. Nagy, N. Buchon, and B.P. Lazzaro, unpublished). We can use these flies to monitor pathogen growth in response to diet without any interference from the humoral immune response. There was no difference in the rate of *P. rettgeri* proliferation in ΔAMP10 hosts provided with either low or high sucrose (Fig. 4A), indicating that the typically observed effect of diet on sensitivity to this infection may be a consequence of diet-dependent immune impairment. In contrast, *Serratia marcescens* continued to proliferate faster in ΔAMP10 flies fed high sucrose than in those fed low sucrose (Fig. 4B), indicating an effect of diet beyond direct mediation of the immune system.

**Fig 4.**
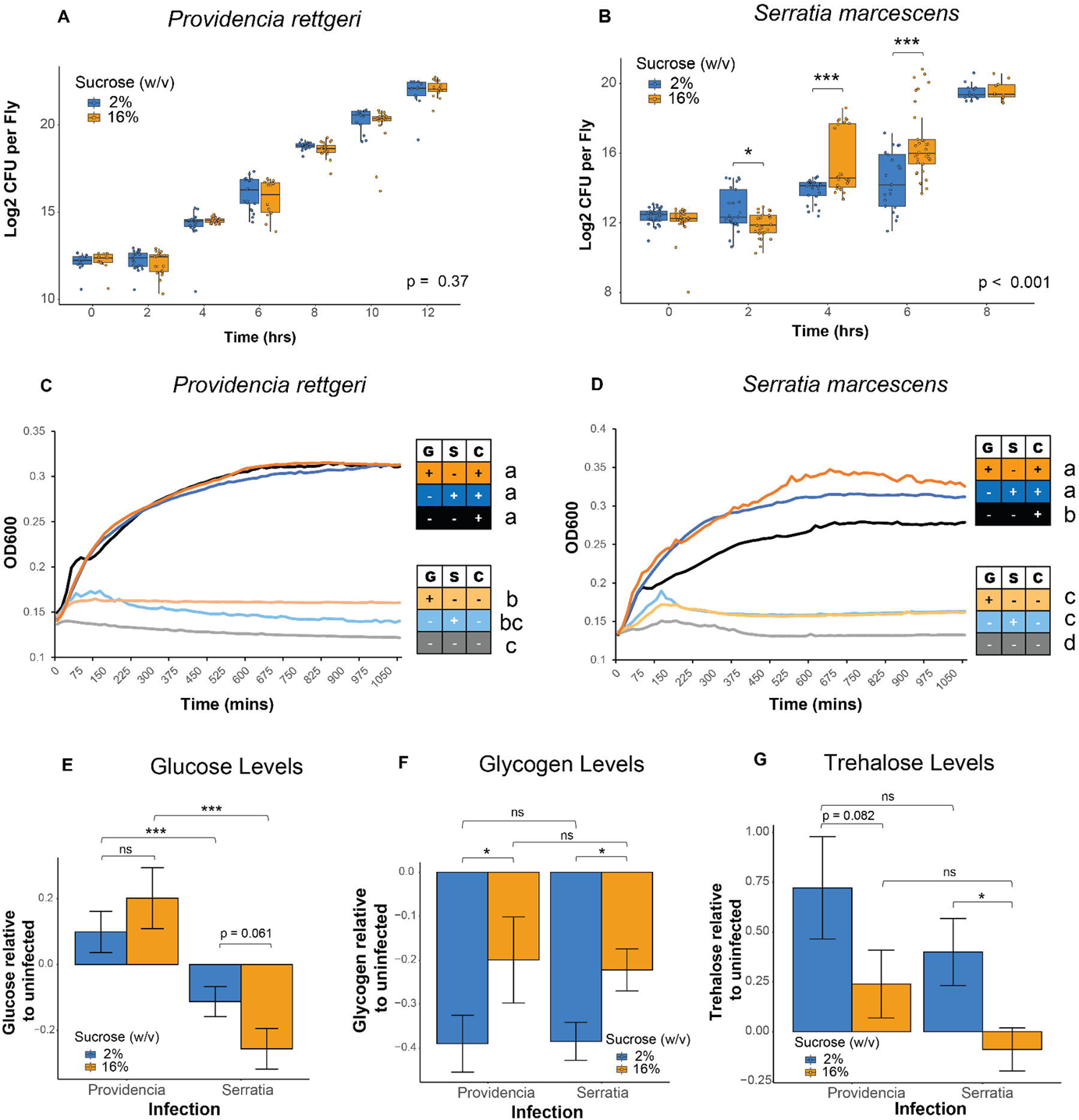
Nutrient availability promotes *Serratia marcescens* proliferation within hosts on high-sugar diets. (A) Diet did not affect the proliferation of *P. rettgeri* in flies lacking 10 major inducible antimicrobial peptides (p> 0.05 time*diet; GLS model). (B) *S. marcescens* proliferate significantly more rapidly in ΔAMP10 flies fed high sugar (p<0.001 time*diet; GLS model). Two hours into the infection, flies fed high sugar exhibit a lower *Serratia* burden (p < 0.01; GLS model followed by Tukey HSD post-hoc), but flies fed high sugar exhibit significantly higher loads by hours by hours 4 (p<0.001) and 6 (p<0.0001; GLS model followed by Tukey HSD-post-hoc). (C) There was no effect of sugar supplementation on *P. rettgeri* growth over 1080 mins in M9 minimal media supplemented with 1% casamino acids (CAS; p> 0.05; ANOVA followed by Tukey HSD post-hoc). CAS supplementation significantly increased OD at 1080 minutes compared to media without casamino acids (p<0.001; ANOVA). (D) *S. marcescens* grew significantly more in minimal media supplemented with the combination of casamino acids and glucose (p<0.001; ANOVA with Tukey HSD post-hoc) or sucrose (p < 0.01, ANOVA with Tukey HSD post-hoc) than in media supplemented with casamino acids alone. *S. marcescens* OD at 1080 mins was significantly lower in media supplemented only with glucose (p < 0.001; ANOVA with Tukey HSD). Each curve represents 6-8 replicates per treatment, pooled. Letters in figures denote significant pairwise differences with p <0.05. Legend: G = glucose, S = sucrose, C = casamino acids. Colors for *in vitro* curves: 1% glucose (light orange), 1% sucrose (light blue), 1% casamino acids (black), or 1% glucose & casamino acids (dark orange), and 1% sucrose & casamino acids (dark blue). (E) Glucose levels declined significantly more after infection with *S. marcescens* but increased in flies infected with *P. rettgeri* on both 2% (p < 0.01; ANOVA followed by Tukey HSD post-hoc) and 16% (p< 0.0001; ANOVA followed by Tukey HSD post-hoc) sucrose diets. There was no significant difference in change in glucose levels after *P. rettgeri* infection between diets (p = 0.24; ANOVA followed by Tukey HSD post-hoc). There was a trend for increased relative loss of glucose after *S. marcescens* infections in flies fed the 16% sucrose diet compared to 2%, however this was not statistically significant (p = 0.0601, ANOVA followed by Tukey HSD post-hoc). (F) Glycogen levels were reduced after infection with both *P. rettgeri* and *S. marcescens*. Flies fed 2% sucrose had significantly higher loss of glycogen stores compared to flies fed 16% sucrose after infection with both *P. rettgeri* (p = 0.0178, ANOVA with Tukey HSD post-hoc) and *S. marcescens* (p = 0.0291, ANOVA with Tukey HSD post-hoc). (G) There was no significant difference in change upon infection in trehalose levels between *P. rettgeri* and *S. marcescens* on either the 2% diet (p = 0.197) or the 16% sucrose diet (p = 0.215). There was a significant reduction in trehalose levels of flies infected with *S. marcescens* on 16% sucrose diets compared to flies fed 2% sucrose (p= 0.0372, ANOVA with Tukey HSD post-hoc). There was a trend toward reduced trehalose in flies provided with 16% sucrose diet after *P. rettgeri* infection, but it was not statistically significant (p = 0.0816, ANOVA with Tukey HSD post-hoc). Legend for panel figure: * = p<0.05, ** = p<0.01, *** = p <0.001.

As an independent, albeit indirect, test of whether *S. marcescens* might utilize excess available carbon to grow faster in hosts provided with high-sugar diets, we measured *in vitro* growth curves of *P. rettgeri* and *S. marcescens* in minimal medium supplemented with either a carbon source or a nitrogen source. We performed an 18-hour growth curve assay in M9 minimal media supplemented with different combinations of casamino acids, sucrose, and glucose. We found that supplementation with 1% casamino acids enabled *P. rettgeri* growth, but there was no further change in the growth trajectory or final OD with supplementation with 1% glucose, 1% sucrose, or no additional sugar (p >0.05, Fig. 4C). These data indicate that *P. rettgeri* is nitrogen-limited but not carbon-limited in minimal media. The addition of either glucose or sucrose dramatically increased the growth of *S. marcescens* in minimal media and resulted in a significantly higher OD at the end of the assay (p<0.01, Fig. 4D), indicating a greater dependence on environmental carbon under these growth conditions. While we do not know precise bioavailability of carbon to bacteria infecting *D. melanogaster*, the data in Figure 2 show that flies on high-sugar diets have higher levels of circulating and stored sugars, and in combination these data offer conceptual support to the hypothesis that the increased proliferation of *S. marcescens* within hosts provided with a high-sugar diet may arise in part from increased availability of carbon to the infecting bacteria.

We had observed that flies provided with high-sucrose diets show elevated glucose and trehalose levels after infection with *S. marcescens* (Fig. 2B, 2C) despite reduction in glycogen stores in both diets (Fig. 2D), leading us to hypothesize that *S. marcescens* might consume excess circulating sugar within the host. We measured the change in carbohydrate levels after infection with either *P. rettgeri* or *S. marcescens* relative to uninfected flies on both diets using the data in Fig. 2B-D. We found that flies infected with *S. marcescens* have a significantly higher loss in glucose levels compared to flies infected with *P. rettgeri* on both the low-sugar (p < 0.01) and high-sugar (p<0.001, Fig. 4E) diets, although they exhibit similar relative loss in glycogen (p > 0.05, Fig. 4F). Interestingly, we observed significantly higher loss of glycogen in flies provided with low-sugar compared to high-sugar diets after both *P. rettgeri* (p = 0.0178) and *S. marcescens* infection (p = 0.029, Fig. 4F). There was no significant difference between the infections in proportional change in trehalose levels on either diet (Fig. 4G). Our observation that both infections stimulate loss in glycogen stores but only *S. marcescens* infection results in a significant reduction in glucose levels is consistent with *S. marcescens* consumption of free sugars during infection.

### *Drosophila* on high sugar diets produce less AMP peptide after infection

We inferred that high-sugar diets increase sensitivity to *P. rettgeri* infection by impairing the immune system because the effect of diet was eliminated in immunocompromised flies. However, the inference that *S. marcescens* may consume excess carbohydrates present in flies on high-sugar diets does not preclude the possibility that those flies might also have reduced immune responses. To directly test whether high-sugar diets impair the immune response, we first measured mRNA transcripts of a panel of genes encoding six *D. melanogaster* antimicrobial peptides (AMPs). Gene expression was measured in flies fed low or high-sugar diets at 8-hours after infection with either *S. marcescens* or *P. rettgeri*. We found that diet had no effect on the infection-induced expression levels of *Diptericin A, Attacin A, Cecropin A1, Drosocin, Drosomycin,* or *Defensin* after either *P. rettgeri* and *S. marcescens* infection (Figs 5A and 5B).

**Fig 5.**
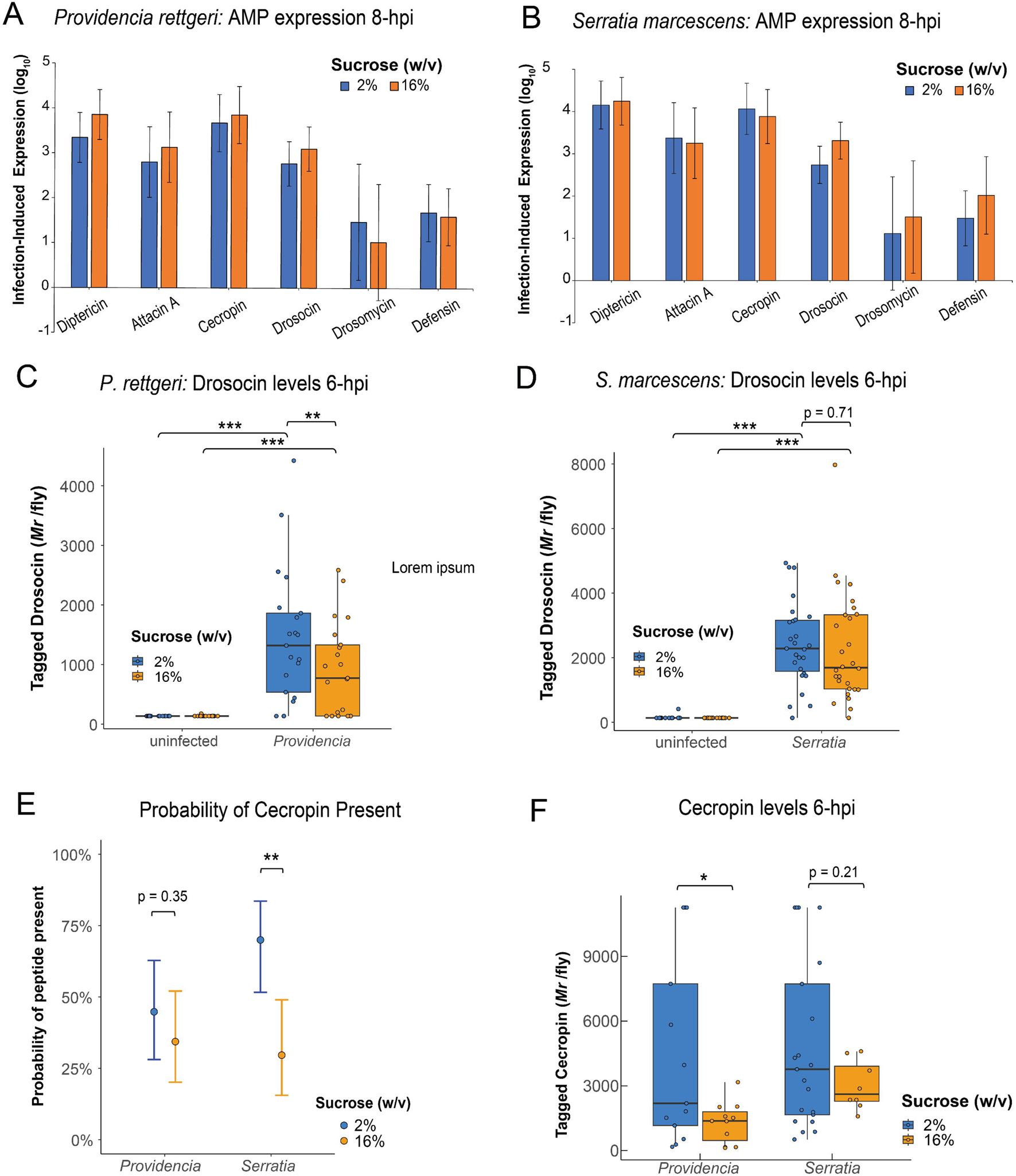
High sugar diets reduce production of AMP peptides in response to infection without affecting AMP gene expression. (A) There was no effect of diet on expression of genes encoding the AMPs Diptericin, Attacin A, Cecropin A1, Drosocin, Drosomycin, and Defensin at 8-hours post-infection with *P. rettgeri* or (B) *S. marcescens* (p > 0.05). (C) *P. rettgeri* infection resulted in significantly higher levels of tagged Drosocin than in uninfected flies on both 2% (p < 0.0001) and 16% (p = 0.0002) diets. Flies on 16% sucrose diet had significantly lower amounts of tagged Drosocin six hours after *P. rettgeri* infection than flies on 2% sucrose diet (p = 0.008). (D) Tagged Drosocin levels were higher at 6 hours after *S. marcescens* infection than in uninfected flies on both 2% (p < 0.001) and 16% (p < 0.001) diets, but there was no difference between the diets after *S. marcescens* infection (p = 0.71). (E) The probability of detectable Cecropin peptide was similar after *P. rettgeri* infection on both diets (p = 0.45), while flies fed the 2% diet were significantly more likely to have detectable peptide present after *S. marcescens* infection than those fed the 16% diet (p = 0.003). (F) Of the flies with detectable Cecropin levels, there were significantly higher peptide levels in flies fed the 2% diet compared to 16% diet after both *P. rettgeri* (p = 0.0229) and *S. marcescens* (p = 0.211) infections.

**Fig S3.**
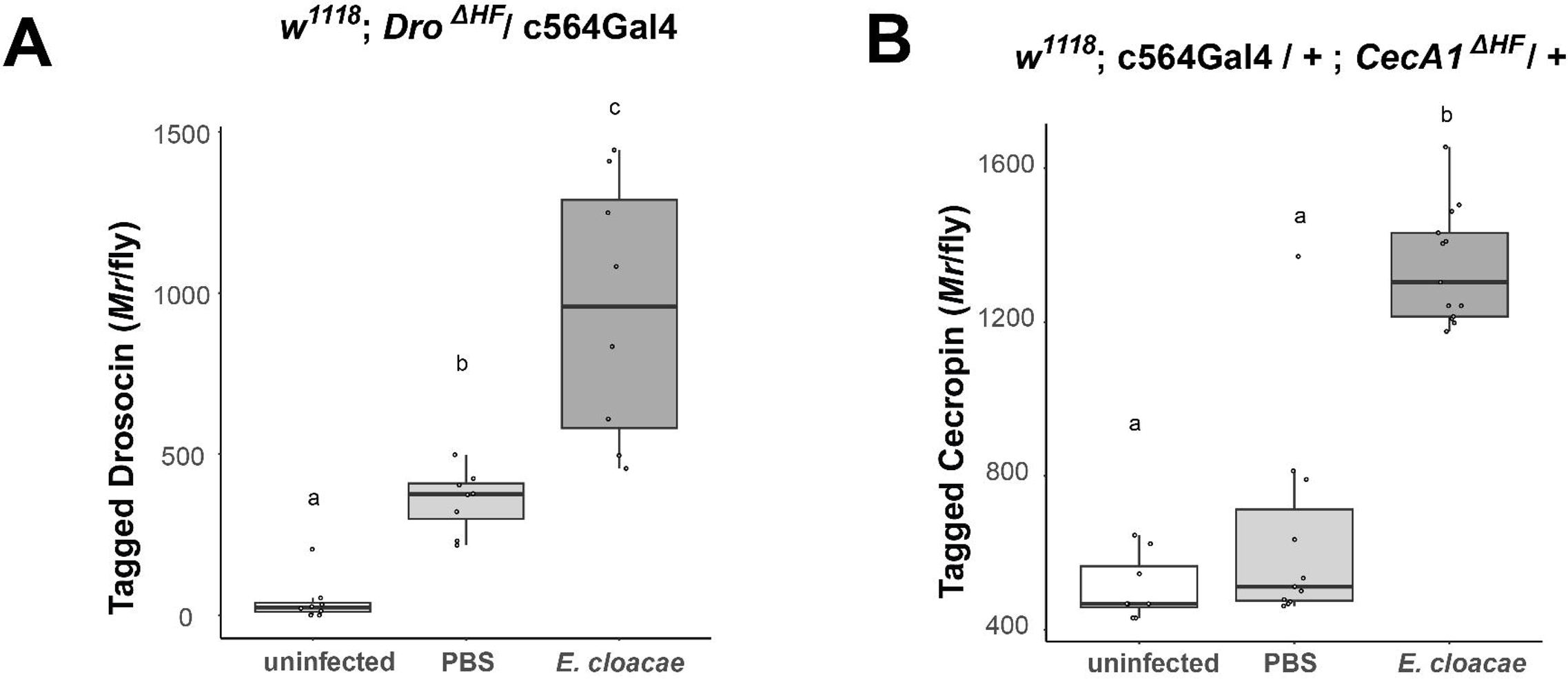
Demonstration that HA-and FLAG-tagged Drosocin and Cecropin can be detected by sandwich ELISA following bacterial infection. (A) Tagged Drosocin levels are significantly higher 3-hours after injection with PBS (p = 0.0455) or ∼6.0×10^4^ cells of the Gram-negative bacterium *Enterobacter cloacae* (p<0.001) than they are in uninjured flies. *E. cloacae* infection results in significantly higher Drosocin levels than PBS injection (p<0.001; linear model with Tukey post-hoc). (B) Tagged Cecropin levels are significantly higher 3-hours after infection with *E. cloacae* than after PBS injection (p<0.001) or in uninjured flies (p<0.001). There was no detectable difference in Cecropin levels between uninjured and PBS-injected flies (p = 0.32). Letters denote pairwise difference p<0.05.

**Fig. S4.**
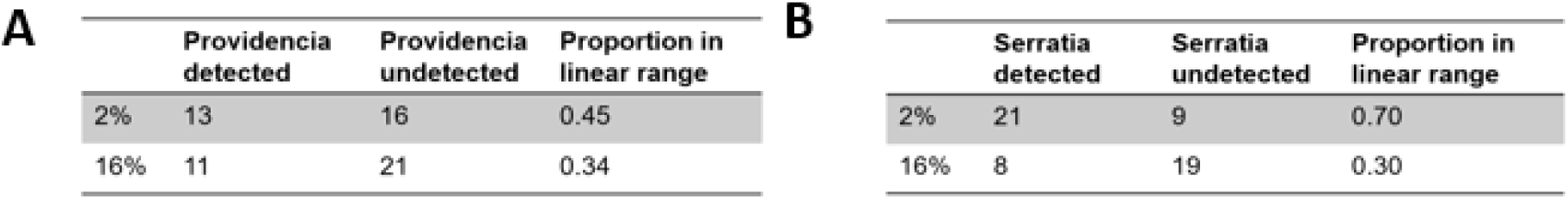
(A) The number of flies fed the 2% and 16% diets that have, or do not have, detectable tagged Cecropin six hours after *P. rettgeri* infection. (B) The number of flies fed the 2% and 16% sucrose diets that have, or do not have, detectable tagged Cecropin six hours after *S. marcescens* infection.

Measuring the amount of mRNA transcript does not directly demonstrate the abundance of translated peptide (57), and translation can be impaired by cellular stress during the *D. melanogaster* response to infection (58). To test whether high-sugar diets impact AMP protein levels in response to infection, we used sandwich ELISA quantify Drosocin and Cecropin A1 from flies that have hemagglutinin (HA) and FLAG epitopes tagged to the endogenous AMPs. Infection-induced production of tagged AMP peptides is detectable as early as 3 hours post-infection (Fig. S3A, S3B). We quantified tagged AMPs from individual flies at 6-hours after infection with either *P. rettgeri* or *S. marcescens.* Flies fed on the high-sugar diet produced significantly lower levels of tagged Drosocin than flies fed on the low-sugar diet after *P. rettgeri* infection (p = 0.008, Fig. 5C). However, we observed no effect of diet on Drosocin levels in flies infected with *S. marcescens* (Fig. 5D). An unexpectedly high number of flies did not have detectable levels of tagged Cecropin peptide after infection with either *P. rettgeri* or *S. marcescens* (Fig. S4A, S4B). We performed a hurdle model to statistically test for differences in Cecropin abundance. The hurdle model first tests the probability that detectable levels of tagged Cecropin are present using a binomial logistic regression, and subsequently tests whether there are quantitative differences in peptide levels conditional on having observed a detectable level of peptide. While the probability of detecting Cecropin was equivalent flies infected with *P. rettgeri* on both diets (Fig. 5E), flies infected with *S. marcescens* had lower probability of expressing detectable Cecropin when fed the high-sugar diet (p < 0.01, Fig. 5E). In the flies for which Cecropin was detectable, peptide abundance was significantly lower in the flies on the high-sugar sucrose diet than in the flies on the low-sugar diet after infection with *P. rettgeri* (p = 0.023, Fig. 5F). Diet had no effect on the abundance of detectable Cecropin after *S. marcescens* infection (p = 0.21, Fig 5F). Overall, flies provided with 16% sucrose diets produce less Drosocin and Cecropin after infection with *P. rettgeri* and have a lower probability of detectable Cecropin 6 hours post-infection with *S. marcescens*. Thus, the translational production of AMPs seems to be impaired in flies on high-sucrose diets even when there is no decrease in transcription level.

## Discussion

In this study, we show that *D. melanogaster* provided with high-sugar diets exhibit higher mortality and elevated pathogen load after infection with the Gram-negative bacteria *Providencia rettgeri* and *Serratia marcescens*, even when the high-sugar diet is provided only to the adult stage after development is complete. *S. marcescens* may be able to take a growth advantage from the excess carbohydrates available in hosts provided with high-sugar diets, and the effect of diet on sensitivity to both infections seems to be mediated by impaired host immune response on high-sugar diet. Interestingly, we saw no effect of diet on susceptibility to infection with the Gram-positive bacteria *Enterococcus faecalis* and *Lactococcus lactis*. As Gram-negative (*S. marcescens* and *P. rettgeri*) infections should primarily activate the IMD signaling pathway whereas Gram-positive (*E. faecalis* and *L. lactis*) infections should primarily activate the Toll pathway (50,59), the much larger effect of diet on sensitivity to Gram-negative infections may implicate an interaction with the IMD pathway.

Our approach of feeding flies on diets that range from 0%-24% sucrose was based on previous studies of systemic infection outcome that defined “high” dietary sugar as ranging from 10% glucose (3) to 24% (w/v) sucrose (12). We predicted that there would be a concentration-responsive decrease in host survival post-infection as dietary sugar increased. However, what we observed was a threshold response where quantitatively variable low-sucrose diets (0%, 2% and 4%) were statistically equivalent to each other but different than high-sucrose diets (16% and 24%) with respect to host survivorship of infection. However, the precise threshold depended on the pathogen, with 8% sucrose being a functionally high-sugar diet for a *P. rettgeri* infection but low-sugar for a *S. marcescens* infection.

Our observation of immunological effects of diet that vary among pathogens is consistent with previous reports in the literature. For example, Ayres and Schneider (2009) (60) demonstrate that dietary restriction at 24 hours prior to infection of adult *D. melanogaster* is protective against a *Salmonella typhimurium* infection, decreases resistance to *Listeria monocytogenes*, and has no effect on *E. faecalis* infection (60). Similarly, Singh et al. (2022) (61) found that *D. melanogaster* reared on a calorie-restricted diet were more susceptible to *Pseudomonas entomophila* infection while diet had no effect on mortality after *E. faecalis* infection. Our finding that diet had no effect on *E. faecalis* infection outcome is corroborated by several studies (60–62) that suggest negligible impact of diet or relevant host physiology on defense against this pathogen. Our data in combination with these previous studies suggest that the effects of diet on immune outcome are in part dependent on the unique physiology of the infecting pathogen.

We observed that *S. marcescens* and *P. rettgeri* pathogen burdens are both higher in *D. melanogaster* fed high sugar. However, the increased pathogen load arises differently for the two pathogens. The effect of diet on defense against *P. rettgeri* is mediated by the host immune system, as flies provided with 16% sucrose diet produce less Drosocin after infection with *P. rettgeri* and the effect of dietary sugar on *P. rettgeri* proliferation is eliminated in *D. melanogaster* lacking a functional immune response. However, while flies infected with *S. marcescens* produced less Cecropin when provided with 16% sucrose diet, we also observed that *S. marcescens* continued to proliferate more rapidly in flies provided with 16% sucrose diets even when those flies lack most inducible AMPs. We found that high-sugar diet increases the level of circulating sugar within the fly, and that *S. marcescens* benefits more than *P. rettgeri* from carbon supplementation when growing *in vitro.* These data suggest *S. marcescens* may perhaps gain a growth benefit from the hyperglycemia of *D. melanogaster* feeding on high-sugar diets. A similar phenomenon has been observed with the Gram-negative opportunistic pathogen *Pseudomonas aeruginosa* infecting hyperglycemic mice. Wild-type *P. aeruginosa* proliferates more quickly in hyperglycemic mice, but mutants for glucose uptake and metabolism genes (*oprB, gltK, gtrS*, *glk)* exhibit similar growth in control and hyperglycemic mice (6). More detailed future study of *S. marcescens* mutants deficient for glucose uptake or metabolism infecting *D. melanogaster* mutant for key factors in metabolism and immune response could probe the relationship between diet, host physiology, and pathogen physiology as determining infection outcome.

Interestingly, we noticed that both wild-type and ΔAMP10 *D. melanogaster* genotypes had lower *S. marcescens* burdens at 2 hours after infection on high-sugar diet than on low-sugar diet (Figs. 3B and 4B). Phagocytosis is known to play a role in controlling *S. marcescens* infection in some contexts (63) and infection has previously been shown to increase aerobic glycolysis in *Drosophila* phagocytes (64). Since ΔAMP10 flies still have intact cellular immune systems, it is possible that the elevated dietary sugar provides energy for phagocytes to use in clearing pathogens sensitive to phagocytosis, although this process alone is not sufficient to manage the infection delivered in our experimental context.

We found that elevated dietary sugar does not impact the transcription of antimicrobial peptides genes (Fig. 5A, 5B), but it does reduce translation of the encoded peptides Drosocin and Cecropin A1 (Fig. 5C-5E). A similar transcriptional effect was observed in larvae of the beet army worm (*Spodoptera exigua*) and fall army worm (*Spodoptera frugiperda*), where infection-induced expression of the AMPs *attacin* and *gloverin* transcripts were equivalent between a low protein and control diet despite reduced resistance to infection on the low-quality diet (65). This suggests that dietary impacts on the immune system may not occur at the transcriptional level but at the level of translation. A recent proteomics study on *D. melanogaster* found that naïve flies subjected to dietary restriction exhibit lower peptide levels of the AMPs Attacin A and Cecropin C (66), although those authors did not measure the effect of diet on production of antimicrobial peptides in the context of a systemic infection.

Antimicrobial peptides are critical for controlling Gram-negative infections in *D. melanogaster*. At the transcriptional level, there is a high degree of co-regulation of genes encoding AMPs (19,67), and we would anticipate corresponding similarity in translational efficiency (68). Given that high dietary sugar reduced production of Drosocin and Cecropin, we anticipate it would also reduce production of other AMPs. It should also be noted that the IMD and Toll immune regulatory pathways do more than simply stimulate production of AMPs. Activation of these pathways alters host metabolism and is influenced by host metabolic status (13), and even stimulates host sequestration of micronutrients to limit pathogens from accessing them (69). Future experiments will be necessary to establish the mechanistic details of how high-sugar diets impact the IMD and Toll pathways to shape resistance to infection.

Our present work establishes that adverse effects of high-sugar diets on infection outcome are due to the combination of impaired host immune function as well as, in some cases, faster proliferation of the infecting pathogen in a hyperglycemic environment. Notably, the immune impairment is not necessarily at the level of transcriptional activation of the immune response but appears to be due to reduced translation of antimicrobial peptides. It is also notable that dietary effects on immune defense are pathogen-specific, and there is not one universal immunological consequence of high-sugar diets. Our findings illustrate the importance of understanding the nutritional requirements of the infecting pathogens as well as the physiological intertwining of host metabolism and immune responses when interpreting effects of diet on infection outcome. We expect this principle to be general across host-pathogen infection systems.

## Methods

### Fly stocks and husbandry

All flies were reared and maintained at 25 °C on a 12-hour light/dark cycle. Mated females of the wild-type strain Canton S (CS) were used for all experiments unless otherwise indicated. Females were cohoused with males in culture chambers and were aged to 3-5 days post-eclosion prior to infection for all experiments. Flies were anesthetized on CO_2_ pads for collection and during infection.

### Generation of Tagged AMP Flies

Flies carrying tagged Cecropin A1 or Drosocin were generated using CRISPR-facilitated homology-directed repair. In brief, guide sequences for *CecA1* or *Dro* were cloned into pCFD5 (*CecA1* guides: guide 1: CCATTGGACAATCGGAAGCT; guide 2: ATAATTATAAATAATCATCG; *Dro* guides: guide 1: CAAAAACGCAAGCAAGCAGC; guide 2: CAATCAATTGTGACACAATG) (70). Homology-directed repair cassettes were cloned into pHD-Scarless-dsRed, encoding the replacement of endogenous *CecA1* and *Dro* with peptides that included FLAG and HA tags (predicted sequences of mature peptides: CECA1^HF^, GWLDYKDDDDKKIGKKIERVGQHTRDATIQGLGIAQQAANVAATYPYDVPDYAR; DRO^HF^, GDYKDDDDKPRPYSPRPTSHPRPIYPYDVPDYARV) (71). These two plasmids were then injected together into embryos of the genotype *y^1^ P(nos-cas9, w-) M(RFP-.attP)ZH-2A w\**. Male flies emerging from this injection were crossed with females of the genotype *w^1118^*; *If /* SM6a or *w^1118^*; TM2 / TM6c, *Sb^1^*, as appropriate. Males expressing dsRed in the eye were then selected and crossed to flies carrying *CyO, Tub-PBac* to enable excision of the 3xP3-dsRed cassette. The final structure of each locus was verified by PCR.

### Experimental Diets

All flies were reared from egg to pupation in rearing chambers with interchangeable food plates on a standard base sucrose-yeast-cornmeal diet (per liter: 60 g yeast, 60 g cornmeal, 40 g sucrose, 7 g *Drosophila* Agar, 0.5 mL phosphoric acid, 5 mL propionic acid, and 6 g methylparaben dissolved in 26.5 mL 100% ethanol). *D. melanogaster* pupated on the chamber walls and, prior to eclosion, the food plates containing the standard rearing media were swapped with one of the experimental diets. Emerging adult flies were aged on the experimental media for 3-5 days post-eclosion prior to infection for all experiments. Experimental diets were identical to the rearing diet aside from the concentration of sucrose. Experimental diets contained either 0%, 2%, 4%, 8%, 16%, and 24% (w/v) of sucrose.

### Infection Survival Assays

Flies were anesthetized on CO_2_ and injected in the thorax with either a bacterial suspension or sterile PBS (as injury control) using a pulled glass capillary needle and nano-injector (72). The following bacteria were injected in at least four experimental blocks: *Providencia rettgeri* strain Dmel, *Serratia marcescens* strain 2698B, *Enterococcus faecalis* strain Dmel*, Lactococcus lactis* strain Dmel. All bacteria were originally isolated as natural infections of *D. melanogaster* (36,73) except *S. marcescens* 2698B, which is a clinical isolate (74). Cultures for *P. rettgeri* and *S. marcescens* were started from a frozen glycerol stock then cultured overnight in liquid lysogeny broth (LB) at 37 °C on a shaker, then subcultured the next morning in fresh LB to achieve growth phase. *L. lactis* was cultured overnight from a frozen glycerol stock in liquid Brain Heart Infusion (BHI) at 37 °C with shaking, then subcultured the next morning in fresh BHI to achieve log-phase. *E. faecalis* was cultured from a frozen glycerol stock at room temperature (∼24 °C) in BHI. All bacteria were resuspended in 1x phosphate-buffered saline (PBS), diluted to A_600_ = 0.1, and injected at a volume of 23 nL per fly. There were ∼4000 *P. rettgeri* cells, ∼3000 *S. marcescens* cells, ∼1000 cells *E. faecalis* cells injected, or ∼ 1000 *L. lactis* cells injected per fly. Flies were housed in groups of ten and each experimental block contained 3-4 vials of flies per experimental diet. Survival curves were assessed using a Cox proportional hazards mixed effect model (coxme) with the “CoxMe” package in R.

Model A: coxme(Surv(status, time) ∼ diet + (1|block)

Diet was treated as a main effect and experimental block was treated as a random factor in our model. This model was used to perform all pairwise comparisons using the “emmeans” package in R with a Tukey p-value correction (p<0.05).

### Bacterial Load Trajectory Assay

To assay bacterial proliferation during *in vivo* infection, flies were infected with *P. rettgeri* or *S. marcescens* as described above, then collected for measurement of bacterial load at 0, 2, 4, 6, 8, 10, 12, 14, 16, 24, 36, and 48 hours post-infection. Single, live flies were homogenized in 500 μL sterile 1x-PBS with a metal bead and 50 μL of the homogenate was plated onto LB agar using a spiral plater (Whitley Automated Spiral Plater-2). To ensure there was no overgrowth of bacteria on the plates, fly homogenates were diluted with PBS at time points where we anticipated higher bacterial growth. For wild-type CS flies, fly homogenates from timepoints 0, 2, and 4 hours were directly plated, homogenates from timepoints 6-12 were diluted 1:10, and homogenates from timepoints 14-48 were diluted 1:100. For *Δ*AMP10 mutants, fly homogenates were directly plated for time points 0, 2, and 4, diluted 1:10 at hour 6, and 1:100 at time points 8, 10 and 12. Agar plates were incubated at 37°C overnight. The resulting colonies were counted using PROTOCOL3 software to estimate the number of colony forming units (CFU) per mL of fly homogenate. We then used the CFU/mL value to approximate the number of bacteria in the individual fly at the time of sampling by using the following formula:

CFU/fly = CFU/mL * (dilution factor) * (0.5 mL/fly of original homogenate)

Two experimental blocks were completed for *P. rettgeri* and *S. marcescens* in CS flies, and three experimental blocks were completed for *Δ*AMP10 mutants with each pathogen. A generalized least squares (GLS) model from the “nlme” package in R was used to determine the effect of diet on pathogen loads over time. Fixed effects in the model include time point sampled (T) and diet (D). The function weights = varIdent(form = ∼1|time point) from the “nlme” package was incorporated into the model to account for unequal variances across time points, which arises as the pathogen grows from the initial injection time point. The emmeans function was used for pairwise comparisons between the experimental diets within each time point using the Tukey method to determine p-values (p<0.05)

Model B: gls(log_2_(pathogen load) ∼ T * D, weights = varIdent(form = ∼ 1|T))

### Excreta Quantification (ExQ) to measure feeding

Methods for the ExQ assay are further detailed in Wu (2020) (47). Briefly, food was prepared by melting down experimental diets and adding 1% (w/v) of dry erioglaucine powder (Sigma # 861146). The dyed food mixture was then dispensed into microcentrifuge tubes caps filled up to 50 μL volume. ExQ chambers were prepared from 50 mL conical vials (Cell Treat # 229421). Air holes were poked along the sides of tubes and the lid using a pushpin (∼0.5 mm diameter). A precision knife was used to cut a hole in the lid of the conical vial to fit the diameter of the food caps. Naïve flies were anesthetized with CO_2_ and sorted into ExQ chambers in groups of 8-10. Every 24 hours, the flies were transferred to new chambers with fresh food. After 48 hours, flies were emptied from chambers and 3 mL of 1x-PBS was used to collect the excreta. Total dye concentration was determined by using a spectrophotometer to measure absorbance of the dye excreted from flies. A standard curve was prepared from a stock solution of erioglaucine by dissolving 10 mg of powder dye in 10 mL of PBS. A serial 2-fold dilution was then created from this initial stock, and the final range of standards used to create the standard curve was 312.5 μg/mL to 5 μg/mL. Absorbance was measured at 630 nm in a 96-well plate on a spectrometer (Molecular Biosciences Spectra Max Series Plus 384). Standards and samples were run in duplicates with 100 μL per well and then averaged for data analysis. Dye consumed per fly within a single ExQ chamber was determined using the following equation:

Dye (μg) per fly = [(Absorbance – Slope)]/(Intercept * # flies)

A two-way Analysis of Variance (ANOVA) in R was used to test for differences in dye consumed by flies on each diet on Day 1 and Day 2 of feeding. Then the emmeans function in R was used for pairwise comparisons between diets within each day using the Tukey test to determine p-values (p<0.05).

Model C: aov(dye ∼ diet + day)

### Quantitative real-time PCR (RT-qPCR) for Antimicrobial Peptide Gene Expression

Female flies provided with 2% and 16% sucrose diets were injected with ∼3000 CFU of log-growth *S. marcescens* or *P. rettgeri* then sampled at 8 hours after injection for quantification of gene expression. Flies were pooled in groups of 5 and frozen at-80 °C for subsequent RNA extraction. In 1.5 mL microcentrifuge tubes, RNA was isolated and purified using a TRIzol extraction procedure according to the manufacturer’s instructions (Zymo Kit direct-Zol RNA kit #R2050). Briefly, flies were homogenized in TRIZoL (Invitrogen #15596018) using a motorized pestle in a microcentrifuge tube and RNA was then extracted using the Zymo Spin Column (Zymo Research #C1078). Samples were then treated with the kit’s DNAse and RNA was dissolved 50 μL of molecular grade water. RNA concentration was determined using a Qubit4 Broad Sense RNA kit (Thermo #Q10210). cDNA was generated using 500 ng of total RNA with the iSynthesis cDNA kit (Bio-Rad #1708891). RT-qPCR was performed on a CFX Connect Real-Time Detection System (Bio-Rad) using PerfeCTa SYBER Green fast mix (VWR #101414-270). Expression of the AMPs *Drosocin, Defensin, CecropinA, Dipterin, Attacin A, Drosomycin* was normalized to the expression of the housekeeping gene *Actin 5C*. Uninfected controls were not subject to injury. Primer sequences used are listed in Supplementary Table 1. A linear model was used to test the contributions of diet and infection status to expression of each AMP gene, standardized to the housekeeping gene (75).

Model D: lm(AMP_CT ∼ *Actin5C_*CT + Diet + Infection Status * Diet)

Diet and Infection Status were fixed effects in the model. Estimated marginal mean CT values for the Diet x Infection interaction terms were extracted using the emmeans package in R. Then we subtracted the estimated marginal mean CT value for uninfected flies from the estimated marginal means from the infected flies for each diet. This difference in estimated marginal mean CT between uninfected and infected values indicates infection-induced change in AMP gene expression on a log_2_ scale. Using the emmeans function, we compared infection-induced expression values between experimental diets with Tukey’s corrections test for p-values (p<0.05).

### Sandwich ELISA to quantify tagged AMPs

HA/FLAG tagged AMPs were quantified 6 hr post-infection. The fly line w^1118^; Dro^HF^/Dro^Hf^ was used to quantify tagged Drosocin and w^1118^; Cec^HF^/+ was used to quantify tagged Cecropin. Maxisorp Nunc 96-well ELISA plates (Thermo Scientific 232698) were coated with 2.5 μg/mL of anti-FLAG antibody (Sigma-Aldrich F1804) in coating buffer (0.2 M sodium carbonate/bicarbonate buffer) overnight at 4 °C, then the plates were blocked for 2 hours at room temperature with 2% bovine serum albumin (Sigma-Aldrich A2153) prior to use. At the time of sampling, flies were homogenized in 200 μL of 0.2% Triton-X buffer and centrifuged at 10,000 RPM for 5 mins to pellet fly debris. Drosocin samples were diluted 1:20 and Cecropin samples were diluted 1:10 so that they fell within linear range of a HA/FLAG peptide standard curve (sequence NH2-DYKDD DDKGG GGGSY PYDVP DYA-NH2 made from AAPTec). The standard curve was prepared by performing a 2-fold serial dilution starting from a stock solution of 2 ng/mL of HA/FLAG peptide to yield 14 standard concentrations. For sample incubation, 50 μL of each standard and sample were added to the ELISA plate and then incubated overnight. Any unused wells were filled with 50 μL of 0.2% Triton-X buffer to serve as blanks. The next day, anti-HA diluted 1:5000 in 0.2% Triton-X was added to each well for another overnight incubation at 4 °C. The following morning, 100 μL of room temperature 1-Step Ultra TMB-ELISA Substrate Solution (ThermoFisher #34028) was added to each well and the plate was incubated at 25 °C for 30 min. After incubation, 100 μL of 2M sulfuric acid was added to each well to stop the reaction. Plates were then read on a spectrometer at 450 nm to measure absorbance. The amount of peptide per fly was determined from the HA/FLAG standard curve with the following equation:

Peptide per fly = [(Absorbance-Slope) * Dilution Factor]/ (Intercept * Fly Homogenate)

To determine whether diet affects Drosocin peptide levels, a linear mixed effects model was applied, with diet and infection status as main factors and experimental block as a random factor. The emmeans function in R was used to compare peptide yield from flies on each diet within infection status.

Model E: lmer(Dro_levels ∼ diet * infection + diet + (1|block))

To determine effects of diet on Cecropin peptide levels, a hurdle model was applied. In the first step, a binomial generalized linear model (GLM) was used to test whether diet and infection status (*Serratia* or *Providencia* infection versus uninjured) affected the probability of detecting Cecropin peptide in the samples.

Model F: glm(Cec_present ∼ diet * infection + diet, family = binomial)

In the second step, a GLM was performed on samples with detectable peptide to test whether diet and infection resulted in quantitative differences in peptide abundance.

Model G: glm(Cec_level ∼ diet *infection + diet)

The significance of the contrast between diets within each infection was determined with a post-hoc Tukey test (p<0.05) using the emmeans package in R.

**Supplementary Table 1:**
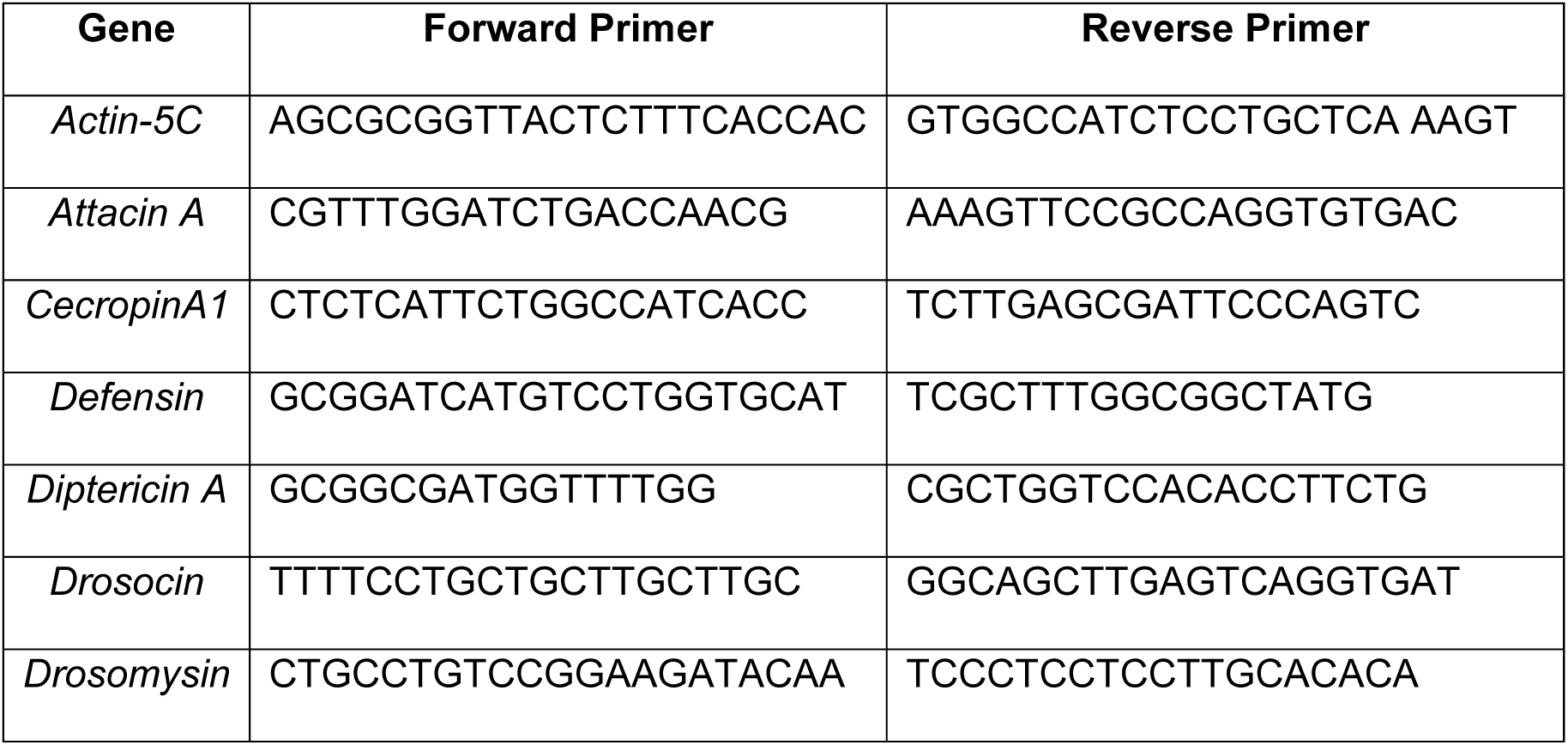
Primer sequences for AMP and housekeeping genes (‘5-3’).

### Measurement of nutritional indices

Nutritional indices were measured in naïve flies and in flies 24 hours post-infection with either *P. rettgeri* or *S. marcescens.* Flies were homogenized in pools of three in 150 μL of cold lysis buffer (10 mM Tris, 1 mM EDTA, pH 8.0 with 0.1% (v/v) Triton-X-100) with a motorized pestle in 1.5 mL microcentrifuge tubes. Homogenates were centrifuged for 1 minute at 13,000 rpm at 4°C to pellet fly debris. The supernatants were transferred to fresh 1.5ml microfuge tubes, and 10 μL of homogenized sample was removed for protein quantification. The remaining samples were placed in a 72°C water bath for 20 minutes to denature endogenous enzymes. The tubes were stored at-80 °C for later use.

### Protein measurements

Protein was measured to normalize all carbohydrate measures. Non-heated fly homogenates were diluted to 1:8 in lysis buffer. Protein was quantified according to Bio-Rad Assay DC Protein Assay Kit instructions. Briefly, 5 μL of standard and sample were incubated with 25 μL reagent A and 200 μL of reagent B. Bovine serum albumin was serially diluted to generate a standard curve that was then used to determine the quantity of protein in each sample. Samples and standards were all run in duplicates. Samples were incubated at room temperature for 15 minutes and then read on a spectrometer plate reader at 750 nm.

### Glucose, trehalose, and glycogen level measurements

Glucose, trehalose, and glycogen levels were measured using reagents from the Sigma Aldrich Glucose (GO) Assay Kit (GAGO20-1KT) and processed according to manufacturer’s instructions. The same fly homogenates were used across all nutrients assayed. Glucose was measured by mixing 5 μL of standard or 5 μL of sample with 150 μL of glucose assay reagent. A glucose standard curve was generated from 2-fold serial dilution from a 1 mg/mL glucose stock concentration. Each sample and standard were run in duplicate. The plate was incubated for 30 minutes at 37°C, then 150 μL of 12M sulfuric acid was added to each well to stop the reaction. Plates were then measured for absorbance at 544 nm on a spectrometer. For trehalose measurements, 5 μL of sample and standards were incubated with 2.5 μL of trehalase overnight at 37 °C. For each sample, two replicates were treated with trehalase enzyme and two were not. Trehalose standard curves were generated from a glucose standard curve, trehalose standards treated with trehalase, and trehalose standards without enzyme. Glucose liberated from trehalose digestion was quantified the next day using the procedure described above, and the abundance of trehalose in the standard was inferred from the difference between the samples that were treated with trehalase and those that were not. Glycogen measurements were conducted similarly, except 5 μL of standard and 5 μL of sample were incubated overnight with amylogucosidase instead of trehalase. The glycogen standard curve was prepared from a glycogen stock solution over the range 1 mg/mL to 0.1 mg/mL. All estimates of carbohydrate levels were normalized to measured protein level for each sample to correct for potential differences in the size of the fly. To calculate change in metabolite level after infection, we subtracted the average post-infection value of the metabolite level from the average level in uninfected samples within an experimental block. A linear mixed-effects model was performed to test for the main effects of diet and infection on carbohydrate/protein ratios with experimental block as a random factor. The emmeans package was used to perform a post-hoc Tukey’s test to determine the significance of pairwise comparisons between diet and infection (p<0.05).

### Bacteria growth curves *in vitro*

*Serratia marcescens* and *Providencia rettgeri* were measured for *in vitro* growth using the kinetic cycle in a spectrometer (Molecular Biosciences Spectra Max Series Plus 384) at room temperature. Bacteria were first cultured overnight with shaking in liquid LB at 37°C. The next morning, bacterial cells were resuspended in 1X M9 minimal media (M9) to achieve A_600_ = 1.0, then further diluted to A_600_ = 0.20 in one of six different experimental M9 media conditions. The combinations of M9 media used were 1% glucose, 1% sucrose, 1% casamino acids (CAS), 1% casamino acids and 1% glucose, 1% casamino acids and 1% sucrose, and a M9 blank. CAS was added to the media to provide a nitrogen source while glucose and sucrose provide a carbon source. In a sterile 96-well plate with a clear flat bottom, 100 μL of bacteria resuspended in the experimental media were pipetted into individual wells in 3-fold replicate. Two wells of uninoculated experimental media were included to monitor for contamination. The plate was covered with a lid and inserted into the spectrometer to measure A_600_ at 15-minute intervals over 18 hours at 37°C. We used a linear mixed effect model to test for the main effects of CAS and sugar with block as a random factor to test whether the addition of CAS and/or sugar led to increased bacterial growth, measured as a higher final A_600_. A post-hoc Tukey HSD was performed using the emmeans package in R to test pairwise comparisons (p<0.05) between media containing sugars and CAS.

Model H: lmer(OD600 at 18 hours ∼ sugar *CAS + sugar + (1|block))

### Preparation of Axenic Flies

Axenic flies were prepared by embryo bleaching. Conventionally reared (microorganism associated) flies were fed on lab standard yeast cornmeal diet (4% sucrose w/v), and approximately 100-150 flies were placed into an egg laying chamber and allowed to lay eggs on grape juice agar plates. Embryos were collected after 20 hours and removed from agar plates with a clean paint brush into a cell strainer. In a six-well tissue culture plate, embryos were washed within the cell strainer first in 10% bleach twice for 1.5 minutes each, followed by two 30-second washes in 70% ethanol. Lastly, embryos were washed twice for 10 seconds with autoclaved 1X PBS. Embryos were continuously mixed throughout the process to prevent them from sticking to each other or the cell strainer. Once sterilized, embryos were transferred with a sterile brush onto autoclaved yeast-cornmeal rearing media in groups of 50. During pupation, the rearing media was replaced with autoclaved 2% and 16% sucrose diets for adult flies to eclose onto.

## Supporting information

Data and Statistics for Fig1-5 and Fig.S1-S3

## Acknowledgements

We thank all members of the Lazzaro lab for feedback on the experimental design for this project and for comments on the manuscript. Figure 1A was designed using biorender.com.

## References

1. Podell BK, Ackart DF, Kirk NM, Eck SP, Bell C, Basaraba RJ. Non-diabetic hyperglycemia exacerbates eisease severity in *Mycobacterium tuberculosis* infected guinea pigs. PLOS ONE. 2012 Oct 4;7(10):e46824.

2. Howick VM, Lazzaro BP. Genotype and diet shape resistance and tolerance across distinct phases of bacterial infection. BMC Evol Biol. 2014 Mar 22;14(1):56.

3. Unckless RL, Rottschaefer SM, Lazzaro BP. The complex contributions of genetics and nutrition to immunity in *Drosophila melanogaster*. PLoS Genet. 2015;11(3):1005030.

4. Kutzer MAM, Armitage SAO. The effect of diet and time after bacterial infection on fecundity, resistance, and tolerance in *Drosophila melanogaster*. Ecology and Evolution. 2016 Jul 1;6(13):4229–42.

5. Cotter SC, Reavey CE, Tummala Y, Randall JL, Holdbrook R, Ponton F, et al. Diet modulates the relationship between immune gene expression and functional immune responses. Insect Biochem Mol Biol. 2019 Jun;109:128–41.

6. SK, Hui K, Farne H, Garnett JP, Baines DL, Moore LS, Holmes AH, Filloux A, Tregoning JS. Increased airway glucose increases airway bacterial load in hyperglycaemia. Sci Rep. 2016 Jun 8;6:27636. doi: 10.1038/srep27636.

7. Thaiss CA, Levy M, Grosheva I, Zheng D, Soffer E, Blacher E, et al. Hyperglycemia drives intestinal barrier dysfunction and risk for enteric infection. Science. 2018 Mar 23;359(6382):1376–83.

8. Gupta S, Koirala J, Khardori R, Khardori N. Infections in diabetes mellitus and hyperglycemia. Infectious Disease Clinics of North America. 2007 Sep 1;21(3):617–38 vii. doi: 10.1016/j.idc.2007.07.003.

9. Lecube A, Pachón G, Petriz J, Hernández C, Simó R. Phagocytic activity is impaired in Type 2 diabetes mellitus and increases after metabolic improvement. PLoS ONE. 2011;6(8).

10. Ilias I, Zabuliene L. Hyperglycemia and the novel Covid-19 infection: Possible pathophysiologic mechanisms. Med Hypotheses. 2020 Jun;139:109699.

11. van Vught LA, Wiewel MA, Klein Klouwenberg PMC, Hoogendijk AJ, Scicluna BP, Ong DSY, et al. Admission hyperglycemia in critically ill sepsis patients: Association With Outcome and Host Response*. Critical Care Medicine. 2016 Jul;44(7):1338–46.

12. Musselman LP, Fink JL, Grant AR, Gatto JA, Tuthill BF, Baranski TJ. A complex relationship between immunity and metabolism in *Drosophila* diet-induced insulin resistance. Molecular and Cellular Biology. 2017 Oct 30;38(2).

13. Darby AM, Lazzaro BP. Interactions between innate immunity and insulin signaling affect resistance to infection in insects. Frontiers in Immunology. 2023;14. Available from: https://www.frontiersin.org/articles/10.3389/fimmu.2023.1276357

14. Buchon N, Silverman N, Cherry S. Immunity in *Drosophila melanogaster* — from microbial recognition to whole-organism physiology. Nature Reviews Immunology. 2014 Dec;14(12):796–810.

15. Bland ML. Regulating metabolism to shape immune function: Lessons from *Drosophila*. Seminars in Cell & Developmental Biology. 2023 Mar 30;138:128–41.

16. Piper MD. Using artificial diets to understand the nutritional physiology of *Drosophila melanogaster*. Current Opinion in Insect Science. 2017 Oct 1;23:104–11.

17. Imler JL, Zheng L. Biology of Toll receptors: lessons from insects and mammals. Journal of Leukocyte Biology. 2004 Jan 1;75(1):18–26.

18. Paquette N, Broemer M, Aggarwal K, Chen L, Husson M, Ertürk-Hasdemir D, et al. Caspase mediated cleavage, IAP binding and ubiquitination: Linking three mechanisms crucial for *Drosophila* NF-κB signaling. Mol Cell. 2010 Jan 29;37(2):172.

19. Troha K, Im JH, Revah J, Lazzaro BP, Buchon N. Comparative transcriptomics reveals CrebA as a novel regulator of infection tolerance in *D. melanogaster*. PLoS pathogens. 2018;14(2):e1006847.

20. Valanne S, Wang JH, Rämet M. The *Drosophila* Toll signaling pathway. The Journal of Immunology. 2011 Jan 15;186(2):649–56.

21. Myllymäki H, Valanne S, Rämet M. The *Drosophila* Imd signaling pathway. The Journal of Immunology. 2014 Apr 15;192(8):3455–62.

22. Tanji T, Hu X, Weber ANR, Ip YT. Toll and IMD pathways synergistically activate an innate immune response in *Drosophila melanogaster*. Mol Cell Biol. 2007 Jun;27(12):4578–88.

23. De Gregorio E. The Toll and Imd pathways are the major regulators of the immune response in *Drosophila*. The EMBO Journal. 2002 Jun 3;21(11):2568–79.

24. Martínez BA, Hoyle RG, Yeudall S, Granade ME, Harris TE, Castle JD, et al. Innate immune signaling in *Drosophila* shifts anabolic lipid metabolism from triglyceride storage to phospholipid synthesis to support immune function. PLOS Genetics. 2020 Nov 23;16(11):e1009192.

25. Dionne MS, Pham LN, Shirasu-Hiza M, Schneider DS. Akt and foxo dysregulation contribute to infection-induced wasting in *Drosophila*. Current Biology. 2006 Oct 24;16(20):1977–85.

26. Chambers MC, Song KH, Schneider DS. *Listeria monocytogenes* infection causes metabolic shifts in *Drosophila melanogaster*. PLOS ONE. 2012 Dec 13;7(12):e50679.

27. Davoodi S, Galenza A, Panteluk A, Deshpande R, Ferguson M, Grewal S, et al. The immune deficiency pathway regulates metabolic homeostasis in *Drosophila*. The Journal of Immunology. 2019 May 1;202(9):2747–59.

28. DiAngelo JR, Bland ML, Bambina S, Cherry S, Birnbaum MJ. The immune response attenuates growth and nutrient storage in *Drosophila* by reducing insulin signaling. Proceedings of the National Academy of Sciences of the United States of America. 2009 Dec 8;106(49):20853–8.

29. Nunes RD, Drummond-Barbosa D. A high-sugar diet, but not obesity, reduces female fertility in *Drosophila melanogaster*. Development. 2023 Oct 16;150(20):dev201769.

30. Hemphill W, Rivera O, Talbert M. RNA-sequencing of *Drosophila melanogaster* head tissue on high-sugar and high-fat diets. G3: Genes, Genomes, Genetics. 2018 Jan 1;8(1):279–90.

31. Ma X, Nan F, Liang H, Shu P, Fan X, Song X, et al. Excessive intake of sugar: An accomplice of inflammation. Frontiers in Immunology [Internet]. 2022 [cited 2023 Apr 25];13. Available from: https://www.frontiersin.org/articles/10.3389/fimmu.2022.988481

32. Yu S, Zhang G, Jin LH. A high-sugar diet affects cellular and humoral immune responses in *Drosophila*. Experimental Cell Research. 2018 Jul;368(2):215–24.

33. Baucom RS, de Roode JC. Ecological immunology and tolerance in plants and animals. Functional Ecology. 2011;25(1):18–28.

34. Musselman LP, Fink JL, Narzinski K, Ramachandran PV, Hathiramani SS, Cagan RL, et al. A high-sugar diet produces obesity and insulin resistance in wild-type *Drosophila*. DMM Disease Models and Mechanisms. 2011 Nov;4(6):842–9.

35. Lazzaro BP, Sackton TB, Clark AG. Genetic variation in *Drosophila melanogaster* resistance to infection: A comparison across bacteria. Genetics. 2006 Nov 1;174(3):1539– 54.

36. Juneja P, Lazzaro BP. *Providencia sneebia* sp. nov. and *Providencia burhodogranariea* sp. nov., isolated from wild *Drosophila melanogaster*. International Journal of Systematic and Evolutionary Microbiology. 2009;59(5):1108–11.

37. Vandehoef C, Molaei M, Karpac J. Dietary adaptation of microbiota in *Drosophila* requires NF-κB-dependent control of the translational regulator 4E-BP. Cell Reports. 2020 Jun 9;31(10):107736.

38. Sharon G, Segal D, Ringo JM, Hefetz A, Zilber-Rosenberg I, Rosenberg E. Commensal bacteria play a role in mating preference of *Drosophila melanogaster*. Proceedings of the National Academy of Sciences. 2010 Nov 16;107(46):20051–6.

39. Fink C, Staubach F, Kuenzel S, Baines JF, Roeder T. Noninvasive analysis of microbiome dynamics in the fruit fly *Drosophila melanogaster*. Applied and Environmental Microbiology. 2013;79(22):6984–8.

40. Shin SC, Kim SH, You H, Kim B, Kim AC, Lee KA, et al. *Drosophila* microbiome modulates host developmental and metabolic homeostasis via insulin signaling. Science. 2011;334(6056):670–4.

41. Wong ACN, Dobson AJ, Douglas AE. Gut microbiota dictates the metabolic response of *Drosophila* to diet. Journal of Experimental Biology. 2014 Jun 1;217(11):1894–901.

42. Fischer CN, Trautman EP, Crawford JM, Stabb EV, Handelsman J, Broderick NA. Metabolite exchange between microbiome members produces compounds that influence Drosophila behavior. Scott K, editor. eLife. 2017 Jan 9;6:e18855.

43. Edgecomb RS, Harth CE, Schneiderman AM. Regulation of feeding behavior in adult *Drosophila melanogaster* varies with feeding regime and nutritional state. Journal of Experimental Biology. 1994 Dec 1;197(1):215–35.

44. Yapici N, Cohn R, Schusterreiter C, Ruta V, Correspondence LBV, Vosshall LB. A taste circuit that regulates ingestion by integrating food and hunger signals in brief article a taste circuit that regulates ingestion by integrating food and hunger signals. 2016 April 21; 165(3) 715–729

45. Ro J, Harvanek ZM, Pletcher SD. FLIC: High-Throughput, continuous analysis of feeding behaviors in *Drosophila*. PLOS ONE. 2014 Jun 30;9(6):e101107.

46. Klepsatel P, Knoblochová D, Girish TN, Dircksen H, Gáliková M. The influence of developmental diet on reproduction and metabolism in *Drosophila*. BMC Evol Biol. 2020 Jul 29;20(1):93.

47. Wu Q, Yu G, Park SJ, Gao Y, Ja WW, Yang M. Excreta quantification (EX-Q) for longitudinal measurements of food intake in *Drosophila*. iScience. 2020 Jan 24;23(1):100776.

48. Bajgar A, Dolezal T. Extracellular adenosine modulates host-pathogen interactions through regulation of systemic metabolism during immune response in *Drosophila*. PLOS Pathogens. 2018 Apr 27;14(4):e1007022.

49. Schwenke RA, Lazzaro BP. Juvenile hormone suppresses resistance to infection in mated female *Drosophila melanogaster*. Current Biology. 2017;27(4).

50. Chambers MC, Jacobson E, Khalil S, Lazzaro BP. Consequences of chronic bacterial infection in *Drosophila melanogaster*. PLOS ONE. 2019 Oct 24;14(10):e0224440.

51. Hanson MA, Dostálová A, Ceroni C, Poidevin M, Kondo S, Lemaitre B. Synergy and remarkable specificity of antimicrobial peptides in vivo using a systematic knockout approach. MacPherson AJ, Garrett WS, Hornef M, Hooper LV, editors. eLife. 2019 Feb 26;8:44341.

52. Rolff J, Schmid-Hempel P. Perspectives on the evolutionary ecology of arthropod antimicrobial peptides. Philosophical Transactions of the Royal Society B: Biological Sciences. 2016 May 26;371(1695):20150297.

53. Bulet P, Hetru C, Dimarcq JL, Hoffmann D. Antimicrobial peptides in insects; structure and function. Developmental & Comparative Immunology. 1999 Jun 1;23(4–5):329–44.

54. Lazzaro BP, Zasloff M, Rolff J. Antimicrobial peptides: Application informed by evolution. Science. 2020 May 1;368(6490). Available from: http://science.sciencemag.org/content/368/6490/eaau5480

55. Kuroda K, Okumura K, Isogai H, Isogai E. The human cathelicidin antimicrobial peptide LL-37 and mimics are potential anticancer drugs. Frontiers in Oncology. 2015;5. Available from: https://www.frontiersin.org/articles/10.3389/fonc.2015.00144

56. Unckless RL, Howick VM, Lazzaro BP. Convergent balancing selection on an antimicrobial peptide in *Drosophila*. Current Biology. 2016;26(2):257–62.

57. Buccitelli C, Selbach M. mRNAs, proteins and the emerging principles of gene expression control. Nat Rev Genet. 2020 Oct;21(10):630–44.

58. Gupta V, Frank AM, Matolka N, Lazzaro BP. Inherent constraints on a polyfunctional tissue lead to a reproduction-immunity tradeoff. BMC Biology. 2022 Jun 2;20(1):127.

59. Sackton TB, Lazzaro BP, Clark AG. Genotype and gene expression associations with immune function in *Drosophila*. PLOS Genetics. 2010 Jan 8;6(1):e1000797.

60. Ayres JS, Schneider DS. The role of anorexia in resistance and tolerance to infections in *Drosophila*. PLoS biology. 2009 Jul;7(7):e1000150.

61. Singh A, Basu AK, Bansal N, Shit B, Hegde T, Prasad NG. Effect of larval diet on adult immune function is contingent upon selection history and host sex in Drosophila melanogaster. bioRxiv; [Preprint] 2022 [cited 2023 Aug 1]. p. 2022.03.03.482770. Available from: https://www.biorxiv.org/content/10.1101/2022.03.03.482770v1

62. Libert S, Chao Y, Zwiener J, Pletcher SD. Realized immune response is enhanced in long-lived puc and chico mutants but is unaffected by dietary restriction. Molecular Immunology. 2008;8.

63. Nehme NT, Liégeois S, Kele B, Giammarinaro P, Pradel E, Hoffmann JA, et al. A model of bacterial intestinal infections in *Drosophila melanogaster*. PLOS Pathogens. 2007 Nov 23;3(11):e173.

64. Krejčová G, Danielová A, Nedbalová P, Kazek M, Strych L, Chawla G, et al. *Drosophila* macrophages switch to aerobic glycolysis to mount effective antibacterial defense. eLife. 2019;8. Available from: https://www.ncbi.nlm.nih.gov/pmc/articles/PMC6867711/

65. Mason CJ, Peiffer M, Felton GW, Hoover K. Host-specific larval lepidopteran mortality to pathogenic *Serratia* mediated by poor diet. Journal of Invertebrate Pathology. 2022 Aug 13;107818.

66. Gao Y, Zhu C, Li K, Cheng X, Du Y, Yang D, et al. Comparative proteomics analysis of dietary restriction in *Drosophila*. PLOS ONE. 2020 Oct 16;15(10):e0240596.

67. Lemaitre B, Reichhart JM, Hoffmann JA. *Drosophila* host defense: Differential induction of antimicrobial peptide genes after infection by various classes of microorganisms. PNAS. 1997 Dec 23;94(26):14614–9.

68. Vasudevan D, Clark NK, Sam J, Cotham VC, Ueberheide B, Marr MT, et al. The GCN2-ATF4 signaling pathway activates 4E-BP to bias mRNA translation and boost antimicrobial peptide synthesis in response to bacterial infection. Cell Rep. 2017 Nov 21;21(8):2039–47.

69. Iatsenko I, Marra A, Boquete JP, Peña J, Lemaitre B. Iron sequestration by transferrin 1 mediates nutritional immunity in *Drosophila melanogaster*. PNAS. 2020 Mar 31;117(13):7317–25.

70. Port F, Bullock SL. Augmenting CRISPR applications in *Drosophila* with tRNA-flanked sgRNAs. Nat Methods. 2016 Oct;13(10):852–4.

71. Bruckner JJ, Zhan H, Gratz SJ, Rao M, Ukken F, Zilberg G, et al. Fife organizes synaptic vesicles and calcium channels for high-probability neurotransmitter release. Journal of Cell Biology. 2016 Dec 20;216(1):231–46.

72. Khalil S, Jacobson E, Chambers MC, Lazzaro BP. Systemic bacterial infection and immune defense phenotypes in *Drosophila melanogaster*. Journal of Visualized Experiments. 2015 May 13;2015(99).

73. Lazzaro BP. A population and quantitative genetic analysis of the Drosophila melanogaster antibacterial immune response. PhD. Thesis, The Pennsylvania State University; 2002. Available from: https://www.proquest.com/docview/305486474/abstract/CF5BBC5BAB284669PQ/1

74. Jensen EC, Amburn DSK, Schlegel AH, Nickerson KW. A collection of Serratia marcescens differing in their insect pathogenicity towards Manduca sexta larvae. bioRxiv. [Preprint] 2020 Jul 29[cited Nov 2023];2020.07.29.226613. Available from: https://www.biorxiv.org/content/10.1101/2020.07.29.226613v1

75. Gordon KE, Wolfner MF, Lazzaro BP. A single mating is sufficient to induce persistent reduction of immune defense in mated female *Drosophila melanogaster*. Journal of Insect Physiology. 2022 Jul 1;140:104414.

